# Elucidating the membrane-binding mechanism of the ALPS-like motif of the Osh4 protein

**DOI:** 10.64898/2026.11.26.769582

**Authors:** Dinara Konakbayeva, Robert J. Allsopp, Madolyn Britt, Sergei Sukharev, Mihaela Mihailescu, Jeffery B. Klauda, Amy J. Karlsson

## Abstract

Transport of lipids within cells is crucial for cellular functions and is facilitated by lipid transport proteins, like the *Saccharomyces cerevisiae* oxysterol-binding homologue (Osh) proteins. Among these, Osh4 is of particular interest, due to its role in lipid trafficking at the endoplasmic reticulum (ER) and trans-Golgi network (TGN). Here, we elucidate the membrane-binding mechanism of the amphipathic lipid packing sensor (ALPS)-like motif of Osh4 using circular dichroism, fluorescence spectroscopy, isothermal titration calorimetry (ITC), and X-ray diffraction (XRD) integrated with molecular dynamics (MD) simulations. Our findings show environment-dependent structural changes of the ALPS peptide, including a preference for α-helical structure in the presence of phospholipid membranes which is modestly enhanced in the presence of anionic lipids. Fluorescence spectroscopy and MD simulation results corroborate the ALPS peptide’s interaction with model membranes and reveal slightly enhanced binding to membranes containing anionic lipids. Moreover, XRD demonstrated the peptide influences membrane structure, with thinning observed upon peptide incorporation. We determined the binding affinity of the ALPS peptide to model membranes using ITC, which supports enthalpy-driven contributions between anionic lipids and the lysine in the ALPS peptide and an entropic contribution from serine and threonine residues desolvating into the membrane. Comparison with computational estimates using potential of mean force calculations suggests current force fields overestimate binding energy. Overall, our results support that the ALPS-like motif will preferentially bind to the TGN cytosolic leaflet, which contains a higher concentration of anionic lipids compared to the ER, providing insight into lipid transport by Osh4.

## 1. Introduction

The movement of lipids between organelles is essential for ensuring the proper function of the entire cell. Lipids, much like other biomolecules, are predominantly synthesized within the endoplasmic reticulum (ER) before being distributed to various organelles. The internal trafficking of lipids encompasses two principal routes: vesicular and non-vesicular mechanisms. Certain cellular compartments that are not served by vesicular routes necessitate an alternative mode of lipid delivery, highlighting the importance of non-vesicular transport [1]. Even among organelles engaged in the secretory pathway, a significant proportion of lipid transfer occurs through non-vesicular means. Most non-vesicular lipid transfer likely occurs at regions of close apposition between two organelles, commonly known as membrane contact sites (MCSs). Lipid transfer proteins (LTPs) are believed to predominantly facilitate this lipid exchange at MCSs [2, 3].

LTPs represent a peripheral membrane protein subgroup that binds to a membrane’s surface. These proteins are characterized by having binding motifs, such as helical and amphipathic motifs. Others interact with membranes through a mix of non-specific and specific interactions [4]. LTPs form a vast and structurally varied class of proteins that includes members from numerous protein (super)families, including steroidogenic acute regulatory protein–related lipid transfer (StARkin) proteins, oxysterol-binding proteins (OSBPs), bridge-like proteins, prokaryotic proteins, and tubular lipid-binding proteins, among others [1]. The OSBP-related proteins (ORPs) are a large family of LTPs that are conserved across species from *Saccharomyces cerevisiae* to humans [5]. ORPs play crucial roles in a variety of cellular functions, including signaling, vesicular trafficking, lipid metabolism, and the non-vesicular transfer of sterols, phosphatidylserine (PS), and phosphatidylinositol 4-phosphate (PI(4)P). Within the ORP family, the most extensively studied are the mammalian OSBPs and the *S. cerevisiae* oxysterol-binding homologue (Osh) proteins [6]. Humans possess 12 ORP genes, which give rise to 16 distinct protein variants through alternative splicing [5]. In contrast, *S. cerevisiae* has seven ORP genes, known as Osh1 through Osh7 [7]. Osh proteins transport sterols, oxysterols, or PS [8–11], by exploiting the metabolism of the important signaling lipid PI(4)P via coupled exchange of sterols or PS [11–13]. Additionally, Osh4 may use its sterol/PI(4)P exchange activity to regulate PI(4)P content of post-Golgi vesicles, thereby enabling their docking and fusion with the plasma membrane.[14, 15] This discovery expands our understanding of the functional versatility and importance of these proteins within cellular processes.

The idea that MCSs are fundamental to lipid transport has been scrutinized by some researchers who doubt that MCSs are necessary [16]. However, our recent work has shown the potential mechanism and structure in which an LTP, specifically oxysterol-binding homologue 4 (Osh4), adjusts its tertiary structure to form contact between two opposing organelles. This computational work suggests a mechanism for a distinctive dual-membrane-bound state of Osh4 that could stabilize and form its own MCS [17]. To more definitively determine whether membrane contact sites exist, it is necessary to study how protein motifs bind to membrane organelles and whether proteins can bind two membranes simultaneously.

LTPs can have motifs dedicated to sensing particular lipids or membrane characteristics. Lipid packing and curvature, for example, can serve as a signal to LTPs [18, 19]. A good example of such lipid sensors is ArfGAP1, a Golgi-localized protein, which has an amphipathic lipid packing sensor (ALPS) motif that recognizes highly curved membranes through hydrophobic interactions. [20] Similar ALPS motifs have been found in other membrane-binding proteins. [21] ALPS motifs are a specific type of amphipathic helix and are characterized by a high abundance of serine and large hydrophobic residues, as well as a low density of charged amino acids [22, 23]. The Osh4 protein possesses an ALPS-like motif composed of 20-27 amino acids [4, 18, 24]. This study focuses on the sequence that starts at residue 1 on the α-helical N terminus and ends at residue 20 of the Osh4 protein (NH_2_-MSQYASSSSWTSFLKSIASF). This N-terminal sequence (**Figure 1A**), represents one of its main membrane-binding regions [25]. Notably, six residues, Y4, W9, F12, L13, I16, and F19, cluster on one face of the helix to form a predominantly hydrophobic surface that is thought to mediate membrane insertion and curvature sensing.

**Figure 1.**
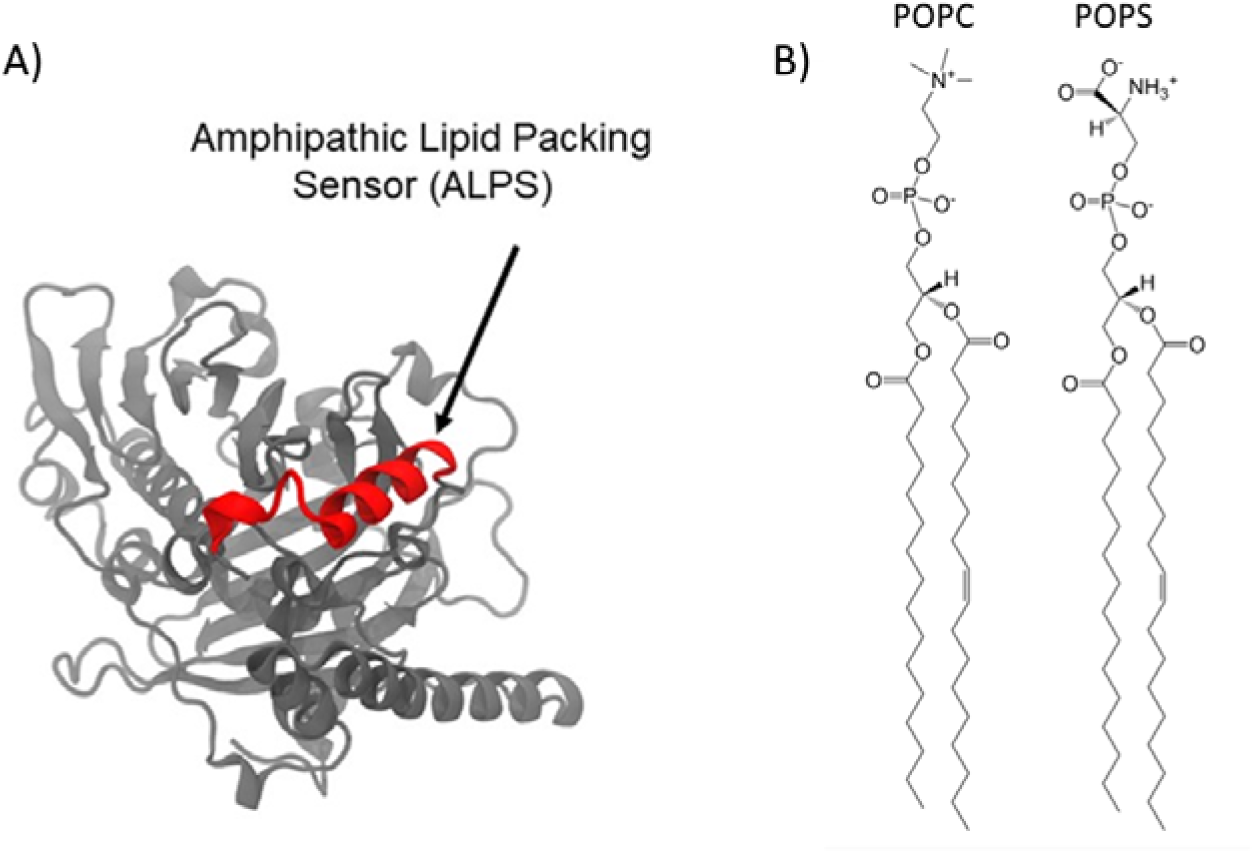
(**A**) Structure of Osh4 with ALPS highlighted in red. (**B**) Structure of 1-palmitoyl-2-oleoyl-glycerol-3-phosphocholine (POPC) and 1-palmitoyl-2-oleoyl-sn-glycero-3-phospho-L-serine (POPS) lipids.

ALPS motifs play a role in cellular stress response and are involved in membrane targeting for lipid trafficking [26]. These motifs prefer binding to membranes that have significant packing defects [26, 27]. The number of packing defects for 1-palmitoyl-2-oleoyl-glycero-3-phosphocholine (POPC) and 1-palmitoyl-2-oleoyl-sn-glycero-3-phospho-L-serine (POPS) is greater than for saturated lipids but less than for lipids with polyunsaturated tails (not native to yeast). Thus, POPC and POPS can provide a representative model for packing in the ER and studying ALPS binding[28]. Due to the abundant presence of uncharged polar residues, such as serine and threonine, on the polar face of the peptide, the ALPS-like peptide exhibits a reduced sensitivity to the typical electrostatic interactions that occur between the predominantly cationic peripheral proteins and the anionic membrane surface [18]. This raises an intriguing question about the necessity of anionic lipids, such as POPS, and zwitterionic phospholipids, such as POPC, for ALPS binding (**Figure 1B**). POPC is a predominant component of cell membranes, and POPS, characterized by its anionic headgroup, is vital in cellular signaling of C-reactive protein and for apoptosis [29, 30].

Anionic lipid interactions are impacted by multiple factors, including overall physiological lipid compositions and the degree of concentration asymmetry across leaflets of the ER, Golgi, and trans-Golgi network. For instance, the lipid distribution of the ER is mostly symmetric between leaflets [31]. The ER membranes are enriched in PC and PE with minority contributions of other lipids. Notably, small concentrations of sphingolipids and cholesterol (or ergosterol in yeast) impact certain membrane properties, such as leading to thinner, looser packing and greater disorder [26, 32]. The ER is also characterized by low anionic charge [26]. The Golgi does have lipid asymmetry because the PE and PS lipids transport to the cytosolic face while PC and sphingomyelin (or inositol-ceramides in yeast) remain on the intracellular face [31]. The Golgi is enriched in PC, PI, and PS, which constitute more than 90% of the top-50 enriched lipids in the Golgi fraction [33]. The trans-Golgi network also has a highly dynamic lipid composition and an asymmetric lipid distribution between leaflets [34]. PS is symmetrically distributed in the yeast ER and Golgi [31, 35, 36], and the lipid compositions in yeast are known for the ER [37], TGN-endosome system [38], and whole cell extract [38]. PS distribution relies on active lipid transfer processes notably mediated by Osh proteins [39, 40].

POPS and other anionic lipids play significant roles in cell biology, and several systematic studies have focused on the role of anionic lipids in interactions with membrane-binding motifs. For instance, the *S. cerevisiae* protein Spo20 has a membrane sensor region that is distinct from the ALPS motif. The Spo20 membrane sensor is sensitive to small variations in anionic lipids and preferentially localizes proteins to the plasma membrane, while the ALPS motif localizes them in the Golgi apparatus [41]. Another study identified an amphipathic helix in the yeast protein Opi1 that selectively binds phosphatidic acid lipids over phosphatidylserine and could play a role in membrane biogenesis [42]. Lastly, cytidine triphosphate (CTP):phosphocholine cytidylyltransferase (CCT) regulates PC synthesis, and its lipid-binding domain contains amphipathic helices that sense PC content in membranes to regulate activation of the enzyme and PC synthesis[43].

The interactions of peripheral membrane proteins with anionic lipids are well known. POPS plays a role in recruitment of peripheral membrane proteins like annexin A2 [44], which is involved in vesicle budding. Binding of peripheral proteins to the anionic phosphatidylinositol phosphate (PIP) lipids can result in the formation of microdomains [44], and some proteins have PIP-specific binding motifs like the pleckstrin homology (PH) domain [45, 46]. Specifically for Osh4, anionic lipids are known to provide an anchor for certain domains [46, 47].

Numerous publications already detail the binding processes of short amphipathic peptides and even computational kinetic studies of the binding process [4, 48, 49]. Our research approach involves studying the equilibrium states of the peptide using enhanced computational sampling techniques and then using the potential of mean force (PMF) to probe the binding and unbinding reaction coordinate. Previously, microsecond simulations have been conducted on the Osh ALPS peptide’s interaction with model membranes and focused exclusively on computational analyses to unravel the binding mechanisms [48]. These long simulations probed the binding mechanism and revealed a relationship between lipid packing defects and binding intermediates. If large packing defects were present, the peptide would fully incorporate itself into the membrane hydrophobic core quickly. However, without plentiful packing defects, the peptide would get caught in the outer charged head group region and wait until the head groups parted enough to insert the hydrophobic core into the membrane stochastically. These preliminary studies show the key characteristics of the final bound orientation, with the hydrophobic core fully inserted into the membrane, but comprehensive equilibrium studies of how lipid head groups from POPC and POPS affect the varied conformations and placements of the lowest-energy states are still needed to understand the impact of different amino acids. Therefore, to address this knowledge gap, we combined several experimental techniques with molecular dynamics (MD) simulations. This multi-faceted approach was utilized to investigate the binding behavior of the Osh4 protein’s ALPS peptide to model bilayers of POPC-POPS mixtures. We aimed to enhance our understanding of ALPS motif interactions with membranes and to verify the precision of our computational models.

## 2. Methods

### 4.1. Peptide Synthesis and Preparation

The ALPS peptide (MSQYASSSSWTSFLKSIASF) was commercially synthesized with an amidated C-terminus as an HCl salt (Biosynth) and purified to >95 %. The peptide was dissolved in ultrapure water and dialyzed overnight against water or buffer, as appropriate for the experiment, to remove the HCl salt.

### 4.2. Vesicle Preparation

Large unilamellar vesicles (LUVs) were used to study the binding interaction of ALPS with model membranes. The lipids POPC and POPS in chloroform were obtained from Avanti Polar Lipids, Inc. (USA). Prior to use, chloroform from pure POPC and mixtures of POPC and POPS was removed under a stream of dry nitrogen. The lipids were further dried under a vacuum overnight and then hydrated in water above the transition temperature for 30 min. The suspensions were extruded using Avanti’s mini-extruder through filters with a 0.1 µm pore size to form LUVs. The particle sizes of the resulting vesicles were characterized using the multi-angle dynamic light scattering mode of a Zetasizer Ultra (Malvern Panalytical) and were 110 nm to 140 nm in diameter.

### 4.3. Circular Dichroism

Circular dichroism (CD) spectroscopy experiments were performed to assess the conformation of the ALPS peptide. ALPS samples at a final concentration of 45 µmol/L were prepared in water or trifluoroethanol (TFE) in the absence and in the presence of POPC and POPC-POPS LUVs at a total lipid concentration of 1.125 mmol/L and a peptide-to-lipid molar ratio of 1:25. A Jasco J-1500 CD Spectropolarimeter (Jasco Analytical Instruments, Tokyo, Japan) was used to perform the measurements. Spectra were acquired at 298 K in the wavelength range of 190 nm to 300 nm, with 0.5 nm step resolution, sweep speed of 50 nm/min, a response time of 2 s, and a bandwidth of 1 nm, using a 0.1 cm path length quartz cuvette. The spectra were processed with Jasco software to subtract solvent spectra. The peptide’s secondary structure content was calculated from the CD spectra using Jasco’s Secondary Structure Estimation (SSE) tool. Each experiment was reported as an average of three scans.

### 4.4. X-Ray Diffraction

X-ray diffraction was performed using oriented lipid multilayers with incorporated ALPS peptides. Oriented multilayers were prepared on glass coverslips by incubating 2 mg of a lipid vesicle suspension in water with ALPS peptide (at peptide-to-lipid ratio of 1:25) and then applying the mixture onto glass cover slips. The water was allowed to evaporate slowly, which caused the vesicles to break and fuse into extended bilayer stacks. The samples were equilibrated for at least 4 h before measurements and maintained at 97 % relative humidity using a saturated solution of potassium sulfate (K_2_SO_4_) and a temperature of 298 K during the measurement. X-ray diffraction measurements were performed on a 3-kW Rigaku Smartlab diffractometer. Diffraction spectra were collected in θ*-2*θ (specular) mode by measuring diffracting intensity versus incidence angle (θ). Diffracted intensity for each Bragg order was calculated by integrating the counts below the peaks. The structure factors, denoted as *f(h)*, were determined by calculating the square root of the peak integrals, after subtracting background and applying Lorentz, polarization, beam footprint, and absorption corrections [73]. The electron density, ρ*(z)*,along the bilayer normal direction *z*, was calculated on an arbitrary amplitude scale, according to the following equation [73, 74]:

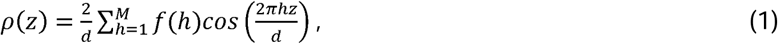

where the *f(h)* is the measured structure factor for the *h*th peak, *d* is the Bragg (repeat) spacing, and *M* is the highest observed diffraction order. ρ*(z)* amplitudes were adjusted using arbitrary scale factors to match at *z*=0 for an easier comparison. Repeat spacings (*d*) and their uncertainties, reported here as one standard deviation, were determined by linear fitting of the Bragg peak positions versus diffraction index (*h*).

### 4.5. Fluorescence Spectroscopy

Fluorescence spectroscopy was used to examine the tryptophan fluorescence emission spectrum of the ALPS peptide in the presence of LUVs. The ALPS peptide was prepared as a 40 µmol/L stock solution, dissolving it either in water or in a buffer solution (50 mmol/L KCl, 9.3 mmol/L K_2_HPO_4_, and 0.7 mmol/L KH_2_PO_4_) at pH 7.0. The LUVs were prepared using the same buffer as the peptide. Titrations were conducted by measuring the emission spectra with the ALPS peptide at a consistent concentration of 20 μmol/L. The lipid concentrations ranged from 0 mmol/L to 4 mmol/L, with measurements taken at 2-fold dilutions. Using a Jasco spectrofluorometer FP-8300, we recorded emission spectra from 300 nm to 500 nm at an excitation wavelength of 280 nm, with a 10 nm excitation (Ex) bandwidth and a 5 nm emission (Em) bandwidth. To reduce fluorescence scattering background, polarizers were set at *Ex_pol_* = 90° and *Em_pol_* = 0°. We calculated corrected fluorescence intensities 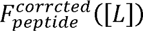 at lipid concentration *[L]* using the tryptophan zwitterion fluorescence as a reference to account for the decrease in the intensity at high lipid concentration associated with losses arising from the scattering of both exciting and fluorescence light [62–64]:

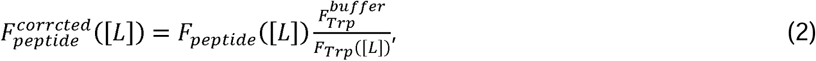

Where *F_peptide_* is the raw uncorrected fluorescence intensity of the ALPS peptide at lipid concentration *[L]*; 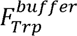 is the fluorescence intensity of pure tryptophan at the same concentration as the ALPS peptide in the absence of LUVs; and *F_Trp_(|L|)* is the fluorescence intensity of pure tryptophan, matched with the concentration of the ALPS peptide in the presence of LUVs at lipid concentration *[L]*.

Fluorescence data were used to predict the dissociation constant *K_d_* using the equation from Tatulian [62–64]:

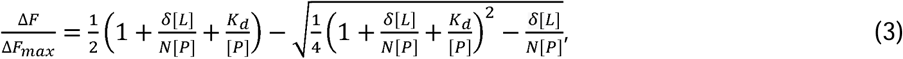

where Δ*F* is fluorescence intensity of the peptide-vesicle mixture minus the fluorescence intensity of the pure peptide, Δ*F_max_* is the saturating value of Δ*F* at high lipid concentration, *[L]* is the lipid concentration, δ is the fraction of the lipid in the external leaflet of the vesicles (δ = 0.52 for 100 nm vesicles), *N* is the number of lipids bound to each peptide molecule, and *[P]* is the total peptide concentration. Theoretical binding curves (Δ*F/*Δ*F_max_* vs *[L]*) were constructed using Equation (3), and the best fit values of *K_d_* and *N* were determined by minimizing the sum-of-squares error. For the binding experiment, two sets of data were collected, averaged, and utilized in the fitting.

### 4.6. Isothermal Titration Calorimetry

Isothermal titration calorimetry (ITC) was utilized to understand the thermodynamics of the binding of the ALPS peptide to vesicles. The ALPS peptide was prepared as a 100-150 µM stock solution by dissolving it in a buffer solution containing 10 mmol/L to 50 mmol/L KCl, 9.3 mmol/L K_2_HPO_4_, and 0.7 mmol/L KH_2_PO_4_ (pH=7). POPC LUVs were prepared at a lipid concentration of 1 mmol/L, and 4:1 POPC-POPS LUVs were prepared at a lipid concentration of 2 mmol/L in the same buffer as the peptides. The LUVs were then dialyzed in the same buffer to minimize the heat of dilution. We injected 10 μL to 15 μL of the LUVs into the reaction cell containing the ALPS peptide with 200 s to 300 s time intervals. The reference power was set to 15 μcal/s, using a syringe stirring speed of 300 rpm. Titrations were carried out on a VP-ITC microcalorimeter (MicroCal, Inc) at 298 K. Three sets of ITC data were collected for each lipid composition. The heat of dilution was not deducted from the reaction heat since it was consistently negligible. The total lipid concentration was used in the estimation of the thermodynamic binding parameters. The experimental data were fit to a one-site model [75]:

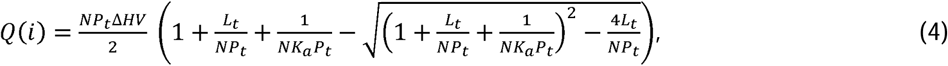

where *Q(i)* the total heat released or absorbed (kcal/mol) for the *i*th injection, N is the number of binding sites, *P_t_*is the total peptide concentration (mmol/L), *L_t_* is the total lipid concentration, *V* is the cell volume (1.32 mL), *K_a_* is the association constant (L/mol), and Δ*H* is the binding enthalpy change. The heat for the *i*th injection, Δ*Q*(*i*), was calculated by subtracting *Q*(*i*-1) from *Q*(*i*) using an equation that includes the correction factor for the displaced volume (i.e., the injection volume *dV_i_*) [75]:

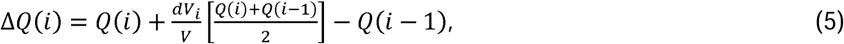

The ITC instrument determines the Δ*Q(i)* for each injection and fits these values to Equation (5) using Origin MicroCal software. The fitting process involves an initial guess of *n*, *K_a_*, and Δ*H*, which are improved by comparing calculated and experimental values. This process is repeated until no further significant improvement in the fit is achieved. The dissociation constant *Kd* is then obtained as the inverse of the association constant (*Kd* = 1/*Ka*).

### 4.7. Molecular Simulations

The membrane models were built with the highly mobile membrane mimetic (HMMM) builder feature within the *Membrane Builder* of CHARMM-GUI (www.charmm-gui.org) [59, 60, 76–80]. These membrane models consisted of POPC and POPS with the ALPS peptide of wild-type Osh4 from the 1ZHY crystal structure [50] with some point mutation modifications to make this the wild-type sequence. The sequence in 1ZHY is **MDPS**QYASSSSW while according to Uniprot the actual sequence is MSQYASSSSW. Additionally, the Protein Data Bank (PDB) file indicates that the first few amino acids are cloning artifacts. The terminal group was patched with the neutral NNEU which is an NH_2_ neutral N-terminus and CT2 patch, which represents an amidated C-terminus to prevent aggregation. The NNEU and CT2 capping method was used to facilitate the comparison with experimental data that would be more prone to aggregate without such modifications. The composition was 4:1 POPC-POPS, with 80 POPC molecules and 20 POPS molecules. The system had 10,000 water molecules and 19 neutralizing potassium ions, calculated from the 20 POPS which are negatively charged and the one lysine amino acid that is positively charged. The amount of water varied depending on the area per lipid, and the TIP3P model [55] was used for water and potassium ions.

The HMMM approach accelerates the binding by enhancing lipid diffusion via replacing the lower part of the lipid tails below the sixth carbon with dichloroethane solvent [59]. The process minimally alters the mechanism of attachment of the peptide onto the membrane [49, 59]. The lipid area scaling factor was varied to achieve simulation box sizes of 74 Å^2^/lipid. These settings help accelerate the binding process by exposing the lipid tails to the solvent and allowing for increased packing defects in the membrane [49, 59]. Each of the models was equilibrated using the standard six-step CHARMM-GUI [77, 78] protocol for 225 ps, as we have reported previously [46]. To simulate the Osh4 ALPS peptide, coordinates from residues 1 to 20 of Osh4 (PDBID: 1zhy) [50] were used. CHARMM-GUI was used to set up the Nanoscale Molecular Dynamics (NAMD) [81, 82] simulations. The process involved minimizing the energy of the system to eliminate unrealistic contacts within the system and the use of restraints in NAMD that were gradually reduced. The equilibration and production runs ran with a 2-fs time step. Other settings for the equilibration included a restart frequency of 500 steps and a trajectory and extended system output frequency of 1000 steps. The peptide was left to simulate and diffuse around the box until it contacted the membrane. The frequency for restart files, dcd (snapshots of the system coordinates used in post processing), xst (system box size information), output energies, and output timing was changed to 5,000 steps for the production runs. All of these HMMM equilibration and production runs used NAMD on a constant number, pressure, area, and temperature (NPAT) ensemble for approximately 150 ns of total simulation time [81, 82]. All simulations used the SHAKE algorithm to constrain hydrogen atoms [83]. Each membrane system had three replicates, each with a unique initial peptide orientation. For all simulations, the CHARMM36 (C36) lipid force field [53] and the C36m additive all atom protein force field [54, 84] parameters were used. All structural equilibrations were performed using the standard CHARMM-GUI approach.

Electrostatic forces and van der Waals forces were computed using a Lennard-Jones force-switching function over 10 Å to 12 Å [85]. The temperature was kept constant at 298 K to ensure a fluid phase membrane using the Langevin thermostat and Langevin dynamics [86]. All simulations were run with periodic boundary conditions (PBC) that were used to evaluate long-range electrostatic interactions using the Particle Mesh Ewald (PME) method [87].

The POPC-POPS lipid simulations with ALPS peptide ran for 300 ns using the last 200 ns of data for the EDP. The control trial ran for 200 ns, and the EDP was based on the last 100 ns of data. The control trial was a mixture of 80% POPC lipids and 20% POPS. This showed that there was not much structural perturbation caused by the presence of the peptide bound to the bilayer of the membrane.

#### HMMM/ Full-Length All Atom (AA)

All HMMM simulations were converted to full-length, all-atom simulations. The conversion required the use of CHARMM-GUI HMMM to full-lipid converter [46]. These conversions were further simulated with NAMD for an additional 300 ns. For the all-atom systems, temperature was held constant as in the HMMM simulations but the box size varied semi-isotropically (*X*=*Y* but not *Z*), while keeping pressure at a value of 1.01325 bar using a Langevin piston [88–90]. This allowed for the membrane to more closely resemble natural membrane lipid packing.

For the analysis of peptide binding to the membranes, the minimum penetration distance graphs were generated by measuring the vertical positions of all the peptide residues in Visual Molecular Dynamics (VMD) [91] around the binding region as well as the average phosphate position over every frame. The figures were rendered with VMD Tachyon internal memory processing [92].

The interaction energy (calculated with the CHARMM program) per residue is the sum of the van der Waals and the electrostatic contributions. CHARMM interaction energy was computed to calculate the energy between the peptide and the membrane, peptide and water, and peptide after it had been removed from the membrane. The interaction energy was calculated for the entire peptide and for each individual amino acid to evaluate the role different amino acids played in the binding of the peptide to the membrane.

#### Potential of Mean Force (PMF)

To obtain free energies of peptide binding to the membrane, the PMF was performed by pulling the peptide off the membrane using an external biasing force. The external bias was applied by sequentially progressing over 1 Å increments to increase the separation distance. After pulling the peptide 1 Å and relaxing for 0.1 ns, the cycle was repeated until the peptide was fully removed from the membrane. Additionally, the membrane was restrained using harmonic restraints to prevent the membrane from stretching or bending to maintain contact with the peptide. The C2 atom of POPC and POPS was selected for restraint because it can be used to divide the head group from the rest of the hydrophobic tail. The C2 atoms of all the lipids were restrained with a constraint scaling factor of 10 kcal/mol/Å^2^ applied to the spring equation. An RMSD collective variable (which is an externally applied force to artificially enforce RMSD) was also applied to the peptide to prevent unfolding and maintain a steady, controlled separation distance. A colvar is an advanced external force that is added into the force field during the MD simulation to maintain a certain distance, or RMSD value. Here we applied a RMSD colvar that maintained the peptide orientation as it was pulled off to avoid unintentional jumps in the PMF profile, as well as to prevent unfolding.

#### Temperature Replica Exchange Molecular Dynamics (T-REMD)

The ALPS peptide was also simulated in a temperature replica exchange molecular dynamics (T-REMD) simulation to quantify the folding free energy landscape. The T-REMD simulations consisted of two replicas for the peptide in 0.15 mol/L KCl ions and four replicas in only neutralizing ions. The number of atoms in these systems was around 43000 atoms to 44000 atoms in total with 14464 molecules of water. The simulations were set up to run 64 replica systems in parallel, each with a slightly different temperature. The temperatures ranged linearly from 293 K to 430 K for all the systems, except one that ranged from 298 K to 450 K. All of the systems were run for 180 ns, and the first 70 ns were discarded as a conservative amount of buffer time for equilibration, leaving 110 ns per simulation of data to be processed. The 0.15 mol/L ion simulation had 2200 frames while the simulation with only neutralizing ions had 4400 frames. About 70 ns were discarded to ensure that the data were fully converged. The free energy of different states was calculated based on binned-state statistics using -*RTln(Probability(x,y)),* where *R*, *T*, and *Probability* are the gas constant, temperature, and probability of a specific state, respectively. The data were processed by creating the counts matrix by discretizing the (*x,y*) data set. Here, (*x,y*) can be the RMSD, radius of gyration, or fraction helicity as defined in the simulation and can be divided by the total number of frames in the simulation to bin the data so that we can apply element-by-element arithmetic in Matlab software using the built in function histcount2 to calculate *-RTln(N_count/total_counts)* where *N_count* is the count of any given bin and *total_counts* is the total number of frames, the free energy comes from the probability of *N_count*/*total_counts*.

#### Electron Density Profiles

The membrane density profiles were constructed by first recentering the bilayer so that the *Z* position of all the atoms was centered. Then, from the centered coordinates, the positions of all the atoms in the system over 200 ns were recorded and a *Z*-based histogram bin for each individual atom in the system was constructed. From there, the data were accumulated for all *Z* positions and all the atoms, and the probability of the atom being found at different *Z* coordinates was determined. Finally, a program called SimtoEXP was used to convert the atomic probability into electron density by multiplying the atoms by the number of electrons they contribute.

#### Calculation of the Simulated *N* Value, Hydrogen Bonds, Peptide Angle, and Contacts Analysis

The MDanalysis methods for amino acid contacts with the membrane are defined by the choline, phosphate, glycerol (O11, C1, HA, HB, C2, HS, O21, C3, HX, HY, O31), C=O (C21, O22, C31, O32), and Cxx which is the first 8 atoms below the unsaturation of the ester. This included the first atom in the double bond on the Sn2 tail of POPC. Each amino acid was selected separately to calculate the contacts. There was a 4.5 Å cutoff for contacts. Then contacts were processed so that within any of the five selections there could only be one contact in each frame. However, the peptide could have contacts with the phosphate and the choline. Then these counts per frame were divided by the total number of frames so that the max contact became 1 and the lowest contact was 0. Then the average of the three trials was taken, and a standard error was calculated to make error bars.

## 3. Results and Discussion

Our experiments and simulations reveal the binding geometry of the ALPS peptide to phospholipid membranes, the structural changes in the membrane upon binding, and the membrane-binding free energy. We focused on membranes composed of POPC and POPS lipids to incorporate lipid-packing defects and varied the POPC:POPS ratio to explore the effect of varied lipid charge. Our membrane models do not conform to the typical concentrations of PS lipids in the ER (2-3% [37]) but represent a model that enhances the concentration of anionic lipids. To probe how the secondary structure of the ALPS peptide changes due to its environment, CD measurements were performed in water, in a hydrophobic solvent, and with LUVs of different lipid compositions. Computational tools, such as aqueous phase replica exchange simulations, were also used to compare with CD results and provide insight into the folding landscape of ALPS in aqueous solution. The binding thermodynamics are essential to probe how lipid composition can influence ALPS binding. To determine the dissociation constant *(K_d_),* we used ITC to obtain binding free energies and compared these with tryptophan fluorescence results fit to a Langmuir adsorption isotherm model. Simulations were also used to obtain estimates for *K_d_* using the PMF along the dissociation reaction coordinate. Finally, membrane structural effects of binding were calculated and measured. We performed X-ray diffraction on multilayer stacks of bilayers to observe the effect of the peptide on membrane thicknesses and compared the results with MD simulations under similar conditions.

### 3.1. ALPS Peptide Structure

Based on the crystal structure for the full-length Osh4 reported by Im et al. [50], the ALPS peptide features a primary helix with distinct polar and nonpolar surfaces (**Figure 2**). To delve deeper into the conformational behavior of the ALPS peptide, particularly during its interaction with membranes, we employed CD spectroscopy. According to our CD results, the ALPS peptide in water is largely unstructured, as reflected by the negative band appearing near 220 nm (**Figure 3**). Estimation of the secondary structure from the CD spectra showed that the majority of the ALPS structure was unstructured, though there were traces of α-helical and β-sheet structures, demonstrating the peptide’s complex and likely dynamic structure (**Table 1**).

**Figure 2.**
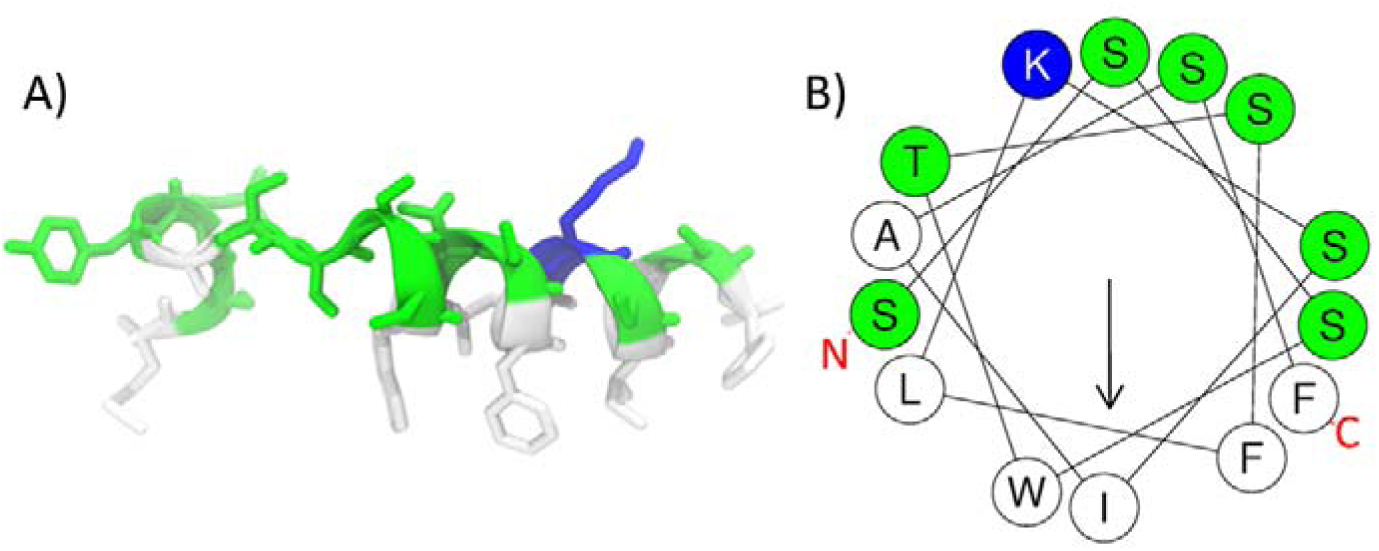
(A) Amphipathic structure of the ALPS peptide. (B) Helical wheel of ALPS that shows where amino acids 7-20 would be located on an α-helix. The helical wheel was created using HELIQUEST.[21] In both panels, hydrophobic residues are color-coded white, polar residues are color-coded green, and the charged residue (+) is highlighted in blue. The arrow in the helical wheel indicates the direction of the hydrophobic moment.

**Figure 3.**
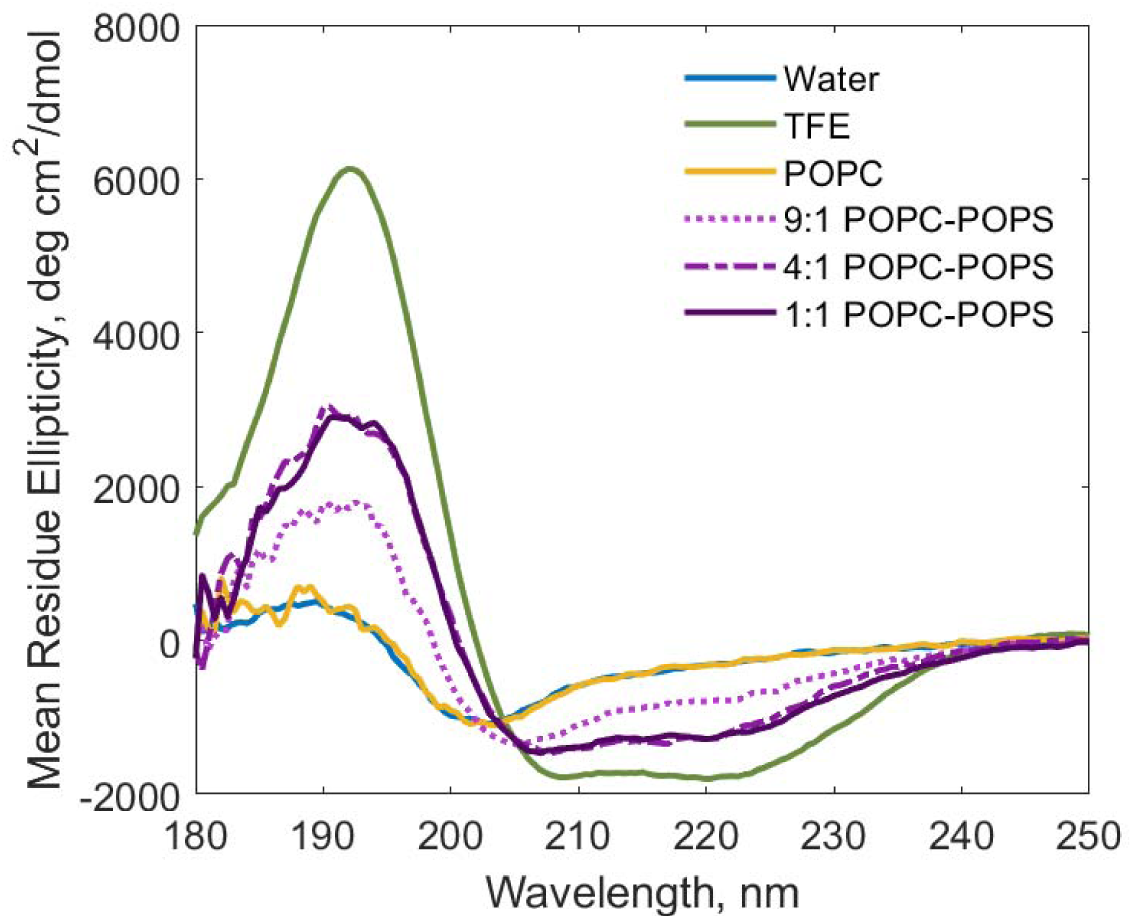
CD spectra of ALPS peptide (45 µM) in the form of mean residue ellipticity measured in water, in TFE, and in the presence of LUVs composed of POPC and POPC-POPS lipids. The final concentration of lipids was 1.125 mmol/L in water. The temperature was maintained at 298 K. Each spectrum is reported as an average of three scans.

**Table 1.**
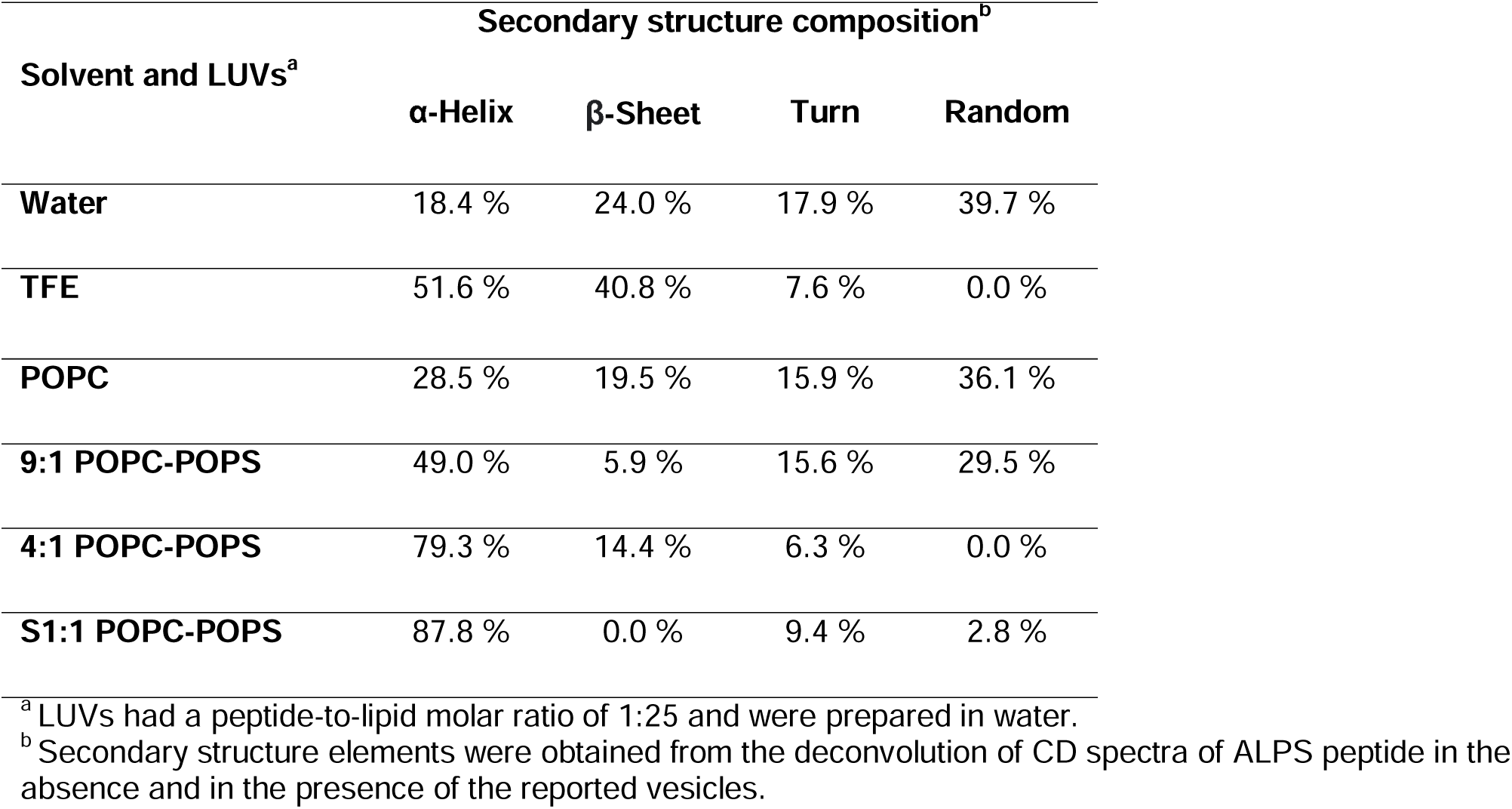
Percentages of secondary structure elements of ALPS in different environments from CD measurements.

Further investigations were conducted using 100 % tTFE, which is a water-soluble alcohol that has amphiphilic character from the combined effect of the trifluoromethyl and the -OH group to mimic the environment of the peripheral phospholipid bilayer [51, 52]. In pure TFE, the ALPS peptide displayed double-minima peaks around 205 nm and 222 nm (**Figure 3**), characteristic of an α-helical structure, with a helical content of 52 % (**Table 1**). Forming a helical structure would allow the hydrophobic regions of the amphipathic ALPS peptide to partition into the hydrophobic environment. Transient polar face contact between the peptide and the membrane would also protect it from exposure to the non-polar solution, thus leading to a lower free energy state.

When we introduced the peptide to large unilamellar vesicles (LUVs) of different lipid compositions, we observed notable changes in its secondary structure compared to the structure in water alone. The helical content increased from 18.4 % in water to 29 % in POPC vesicles (**Table 1**). These results suggest the peptide is slightly more structured in the presence of POPC vesicles than it is in pure water. However, the estimated helical content observed was even higher in TFE than in POPC vesicles.

We also investigated the structure of the ALPS peptide in the presence of vesicles formed from the negatively charged phospholipid POPS. Upon incorporating POPS into the zwitterionic POPC vesicles, the peptide underwent significant changes, showing the double minima (at 210 nm and 222 nm) characteristic of an α-helical structure. The α-helical conformation was more prominent in the presence of PS lipids with 1:1 POPC-POPS (88 %), 4:1 POPC-POPS (79 %), and 9:1 POPC-POPS (49 %) LUVs than in LUVs solely consisting of POPC (29 %). The presence of anionic lipids likely increases the fraction of bound peptide compared to free peptide, and the interaction with the membrane causes folding and increased α-helicity.

Computationally, we used T-REMD to determine how well the protein/lipid force field (CHARMM36m [53]/CHARMM36 [54]) and the TIP3P water model [55] match with the experimental CD data in water. We simulated the ALPS peptide under two conditions: (1) with neutralizing ions and (2) with 0.15 mol/L KCl. Based on the CD experiments, the peptide only has 18 % α-helical character in water, but the T-REMD results suggest the α-helical state is more stabilized in our simulations. For 0.15 mol/L KCl (**Figure 4A**), the T-REMD results show four energy wells at approximately 70%, 55 %, 30 %, and 0 % helical structure. The T-REMD results for neutralizing ions are similar, with folding wells at 75 %, 50 %, and 0 % helical structure (**Figure 4B**). Even though the number of energy wells varied, the average percentage of helical structure of the simulated ALPS peptide with 0.15 mol/L ions was 37 %, which is similar to the percentage helical structure in only neutralizing ions (39 %) but higher than the 18 % helical structure observed in the CD experiments in water.

**Figure 4.**
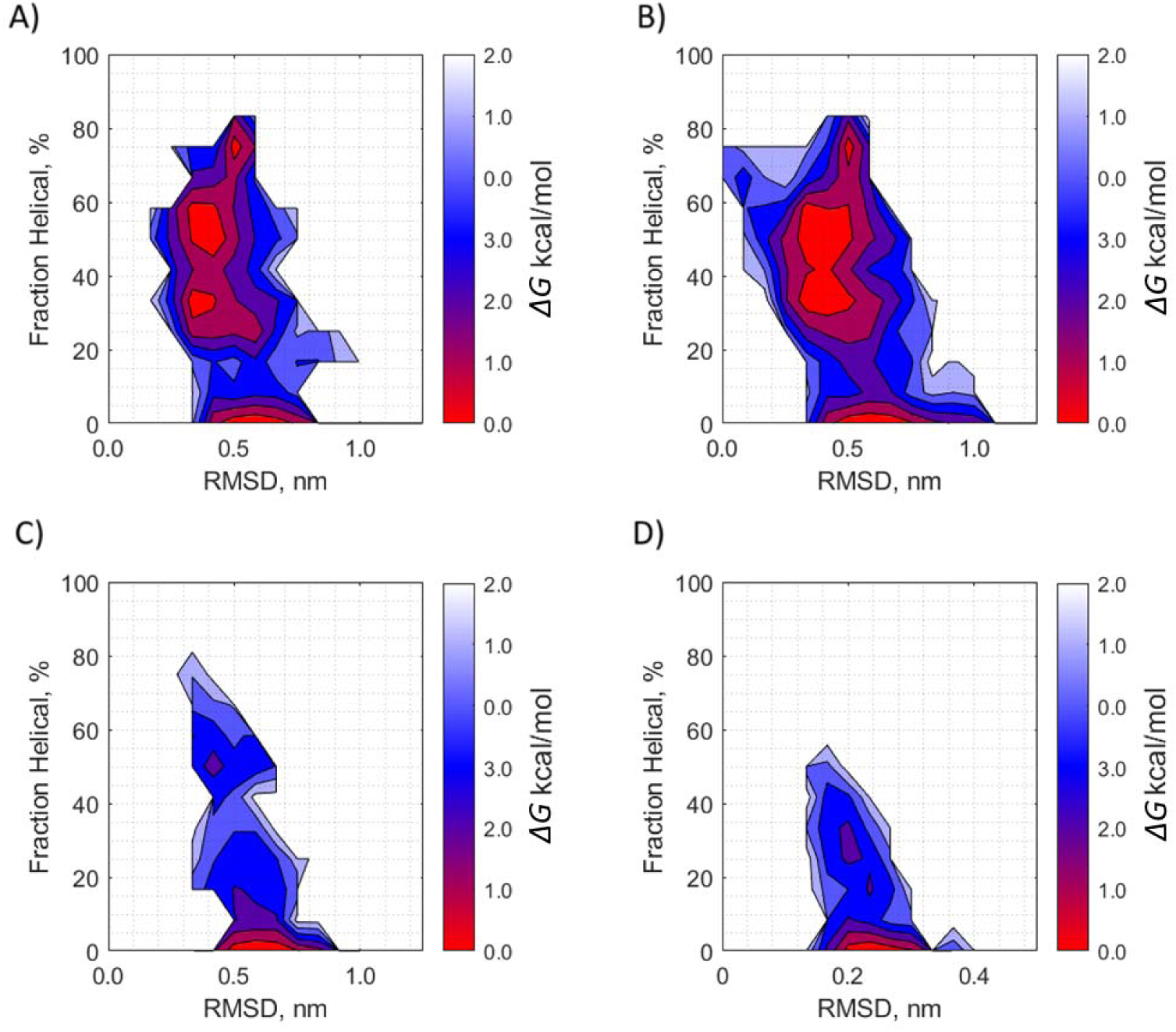
Free energy surface of the ALPS peptide at 298 K in water as a function of fractional helical structure (y-axis) and backbone root mean square deviation (RMSD) (x-axis). Data are shown for the wild-type peptide in (A) 0.15 mol/L KCl and (B) neutralizing ions and for the F13S mutant in (C) 0.15 mol/L KCl and (D) neutralizing ions.

Structures from the different simulations highlight different energy wells, provide a snapshot of the unfolded structure, and illustrate the potential importance of π−π stacking (**Figure 5)**. According to **Figure 5C**, π−π stacking of the W10 and F13 residues takes place in the unfolded state, stabilizing one helical turn. One potential reason for the over-stabilization of the α-helix in the simulations could be an overestimation of the effects of π−π stacking. To test this hypothesis, we simulated the ALPS peptide with an F13S mutation to disrupt the π−π stacking between W10 and F13. For the F13S peptide in 0.15 mol/L KCl (**Figure 4C**) and the F13S peptide in neutralizing ions only (**Figure 4D**), one main energy well is present at low fraction α-helicity, and two to three shallower energy wells are present at higher helicity. Evaluating the distance between the aromatic rings of the W10 and F13 residues (**Figure S1**) shows that the CHARMM36m force field prefers a distance of 5.5 Å for π−π stacking and suggests that the π−π stacking could be a contributing factor in the higher occurrence of folded peptides compared to unfolded in the simulation. The profile for the mutant with disrupted π−π stacking shows up to roughly 15 % helical structure in 0.15 mol/L ions and 10 % helical structure with only neutralizing ions; deep folding wells were only present at 0 % helical structure (**Figures 4C** and **4D**), which better matches the experimental results. Adjustments in the protein force field might be needed to accurately represent the unstructured character for the wild-type ALPS peptide in water.

**Figure 5.**
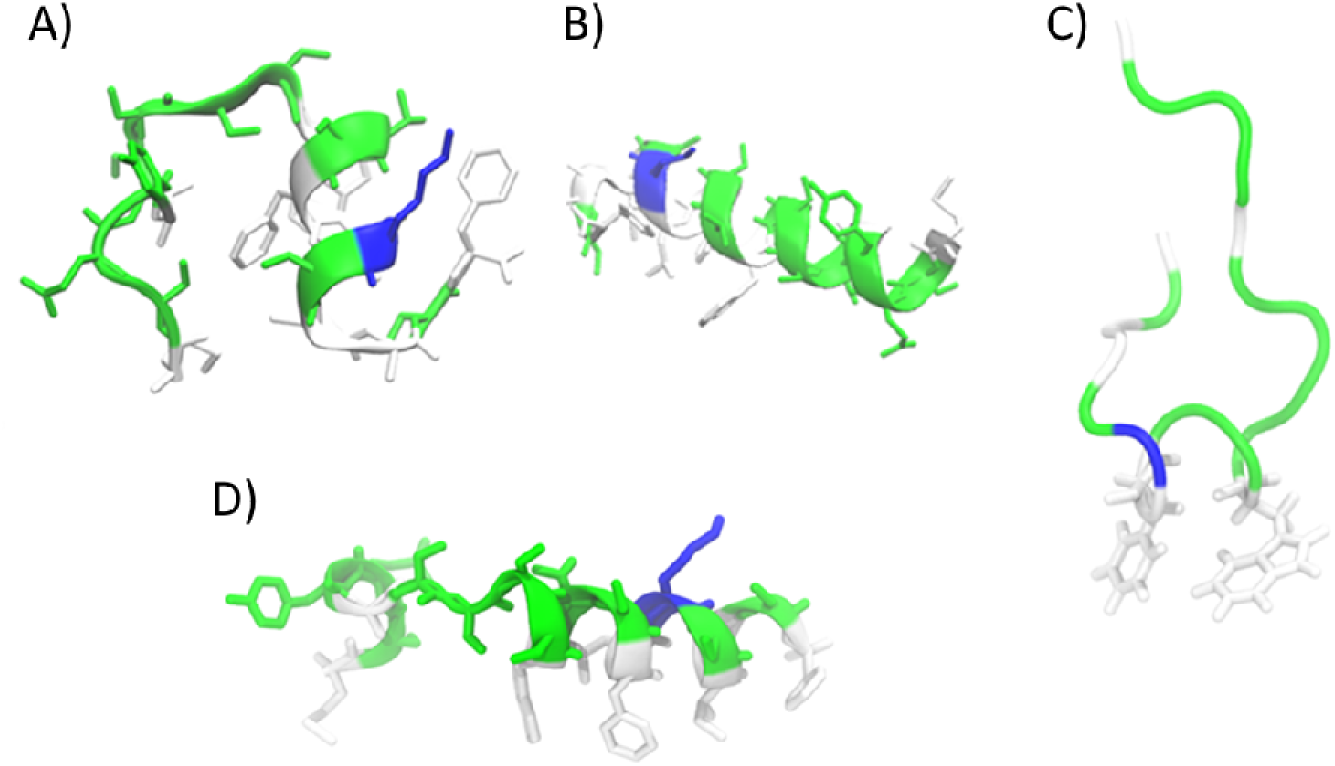
Structures of the Osh4 ALPS peptide in water corresponding to the different folded energy wells. The energy wells are determined by the lowest free energy derived from the probability, which is a function of RMSD (*x)* and helical fraction (*y)*, giving [56]. (A) Structure from simulation with 0.15 M ions that resulted in a helicity of 0.37 and an RMSD relative to the crystal structure of 0.37 nm. (B) Structure from simulation with neutralizing counter ions with a helicity of 0.8 and an RMSD of 0.5 nm. (C) Example structure n the unfolded wells of the mostly unfolded peptide with π−π stacking between W10 and F13 residues. (D) Structure showing a helical conformation with π−π stacking between W10 and F13 residues.

### 3.2. ALPS Membrane Interaction

To investigate how ALPS interacts with model experimental membranes, we first used tryptophan fluorescence spectroscopy to monitor the interaction of the single tryptophan residue in the peptide with LUVs (110 nm to 140 nm, prepared by extrusion). We measured the blue shift of the fluorescence emission maximum of tryptophan due to the residue being in a hydrophobic versus hydrophilic environment [57]. The ALPS peptide in water had a fluorescence emission maximum of 352 nm (**Figure 6**), typical for tryptophan in a polar environment [58]. Upon the addition of POPC-containing LUVs at a concentration of 1 mmol/L, the fluorescence emission maximum of tryptophan shifted toward the blue end of the spectrum (**Figure 6**). The shifts in the maximal wavelength of the spectra suggest that the tryptophan residue of the ALPS peptide is in contact with the hydrocarbon region of the bilayer. Interestingly, when vesicles contained both POPC and POPS lipids, both the blue shift and the maximum tryptophan emission intensity increased. Furthermore, in LUVs with a 4:1 POPC-POPS lipid composition, the blue shift was more pronounced than in those with a 9:1 POPC-POPS ratio. This suggests that the presence of the anionic POPS is facilitating the recruitment of the ALPS peptides to bilayers, likely due to the charged lysine residue present in the ALPS peptide interacting with the negatively charged POPS head group.

**Figure 6.**
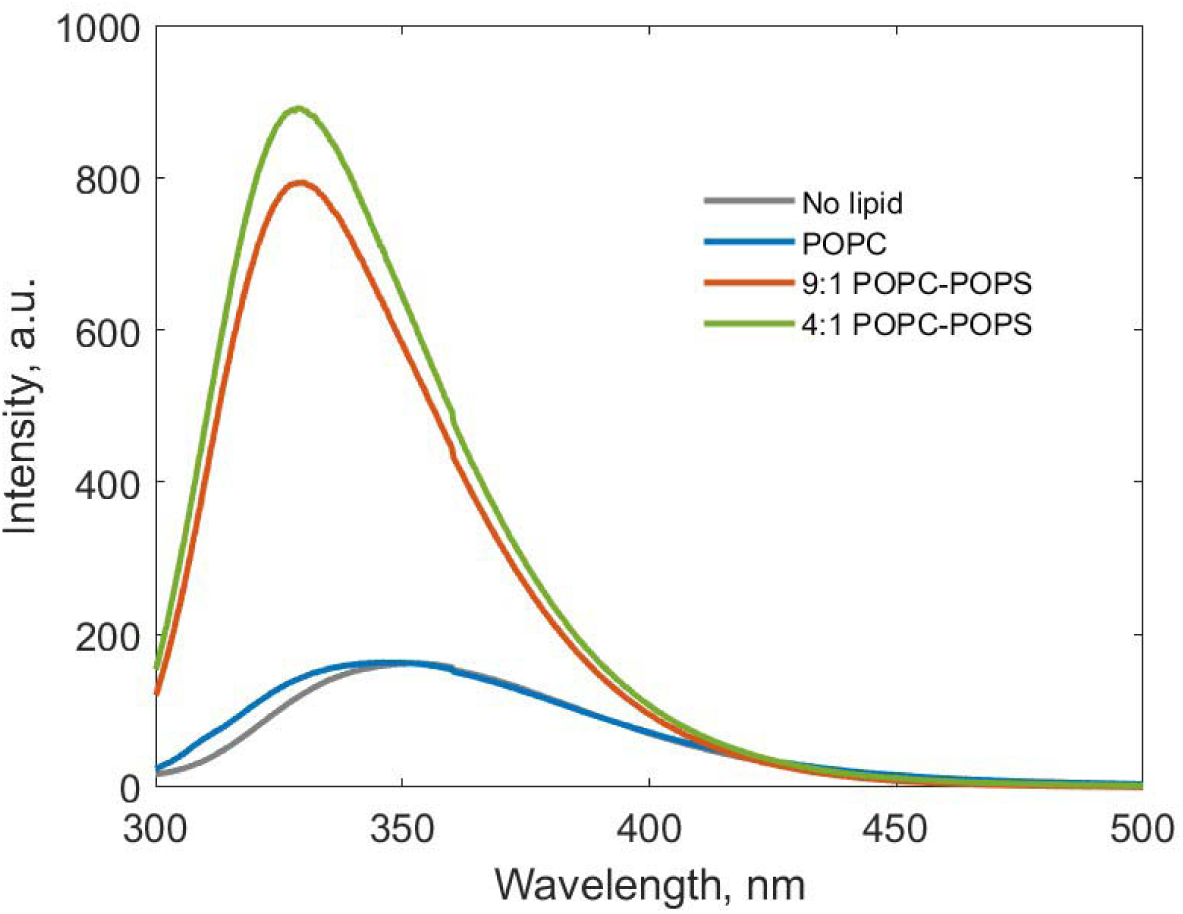
Fluorescence of tryptophan residue of ALPS peptide (20 µmol/L) alone and in the presence of POPC LUVs and POPC-POPS LUVs. LUVs were prepared at three molar ratios of POPC to POPS lipids with total lipid concentrations of 1 mmol/L.

After observing the change in tryptophan environment in the presence of LUVs, we performed MD simulations to probe how the ALPS peptide binds to membranes that correspond to the experimental work. To improve sampling of binding, we used the HMMM approach [59, 60], which replaces lipid tails with dichloroethane to increase head group diffusion and includes an area-per-lipid scaling factor to increase the number of packing defects. The HMMM technique yielded a variety of different ALPS peptide conformations and orientations (**Figures S2** and **S3**). The ALPS tryptophan inserted into the HMMM membrane bilayer, consistent with the tryptophan insertion indicated by the fluorescence spectroscopy experiment results. Using the HMMM technique allowed us to observe variability in the binding and to study the ensemble without being limited by the computationally costly kinetics of binding. To refine the ALPS-membrane structure, full-length lipid simulations were run starting with the HMMM results and converting the reduced membrane to the full-length form. After conversion to the full-length lipids, the simulations were equilibrated for an additional 300 ns. The peptide continued to stochastically adjust the depth of penetration (**Figure S4**) and the angle of the helix formed by the 10^th^ through 20^th^ amino acids (**Figure S5**). The 10^th^ through 20^th^ amino acids formed the most stable α-helix, and this portion laid flat or at various angles and localized in different layers of the membrane at equilibrium (**Figure 7**). **Figure 8** shows, at a higher magnification, the final (equilibrated) structures of ALPS in a POPC-POPS membrane and demonstrates that tryptophan is inserted into the membrane, agreeing with the tryptophan fluorescence data.

**Figure 7.**
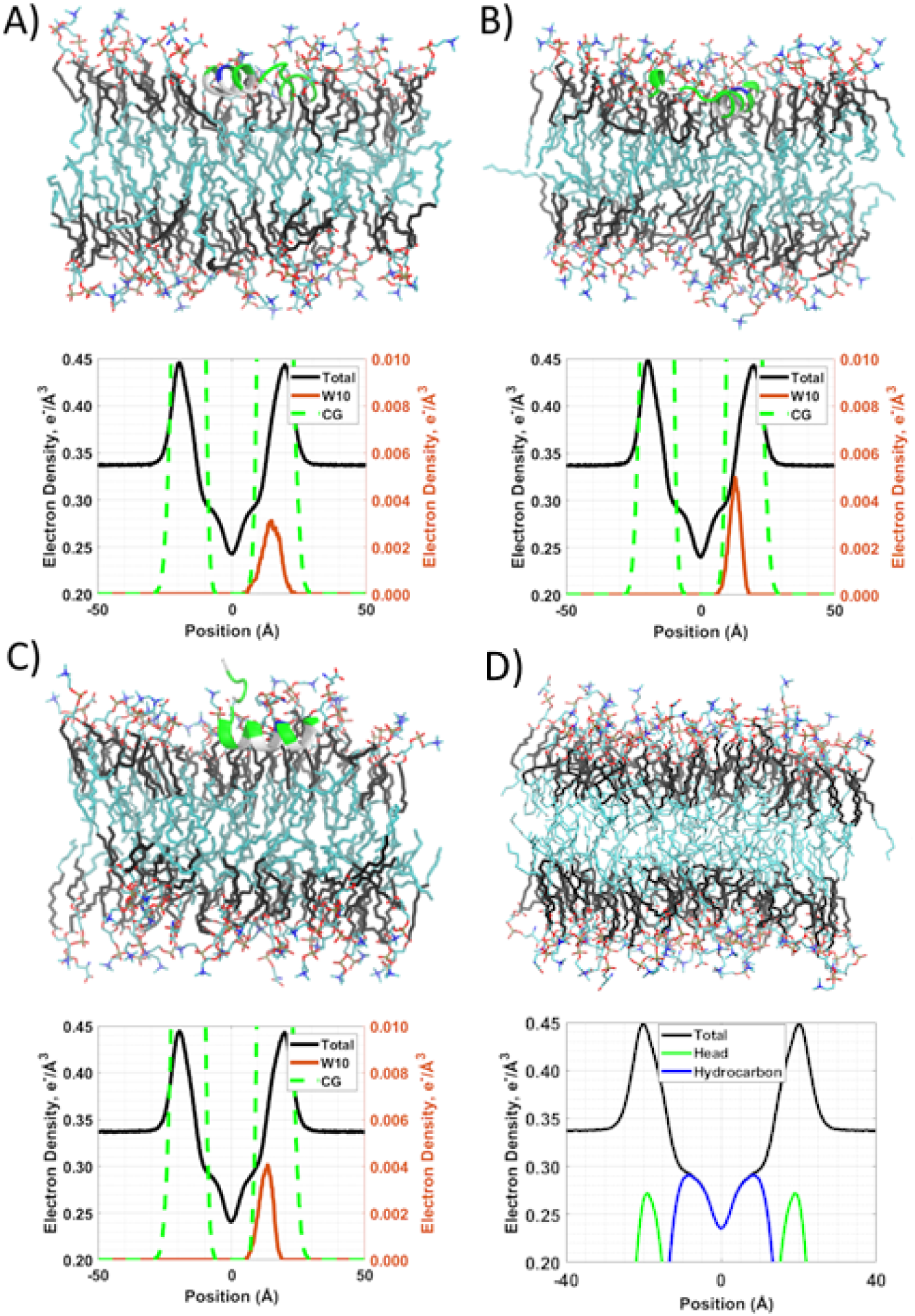
Equilibrated all-atom membrane-peptide system showing the final snapshots and electron density profiles for three replica systems with peptide (A for Replica 1, B for Replica 2, and C for Replica 3) and a control with no peptide D and no POPS to show if either one had any impact on structure. In addition to the total electron density, the density for the carbonyl glycerol (CG) for the replica or the head group for the control, along with the density for the W10 residue, are shown. The density for the W10 and CG residues are shown on a secondary axis on the right of each plot because it is much less intense than the rest of the membrane. The CG group differs from the head group because it is positioned lower down the Z coordinate axis and because of this it is often used as a point of reference in the membrane. The simulations ran for 300 ns. The control trial ran for 200 ns and the EDP was based on the last 100ns of data. The control trial was a mixture of POPC-POPS (4:1) lipids. The lipid is color-coded so that CH_2_ above the double bond is black, CH_x_ below the double bond is cyan, oxygen atoms are red, and phosphate groups are tan.

**Figure 8.**
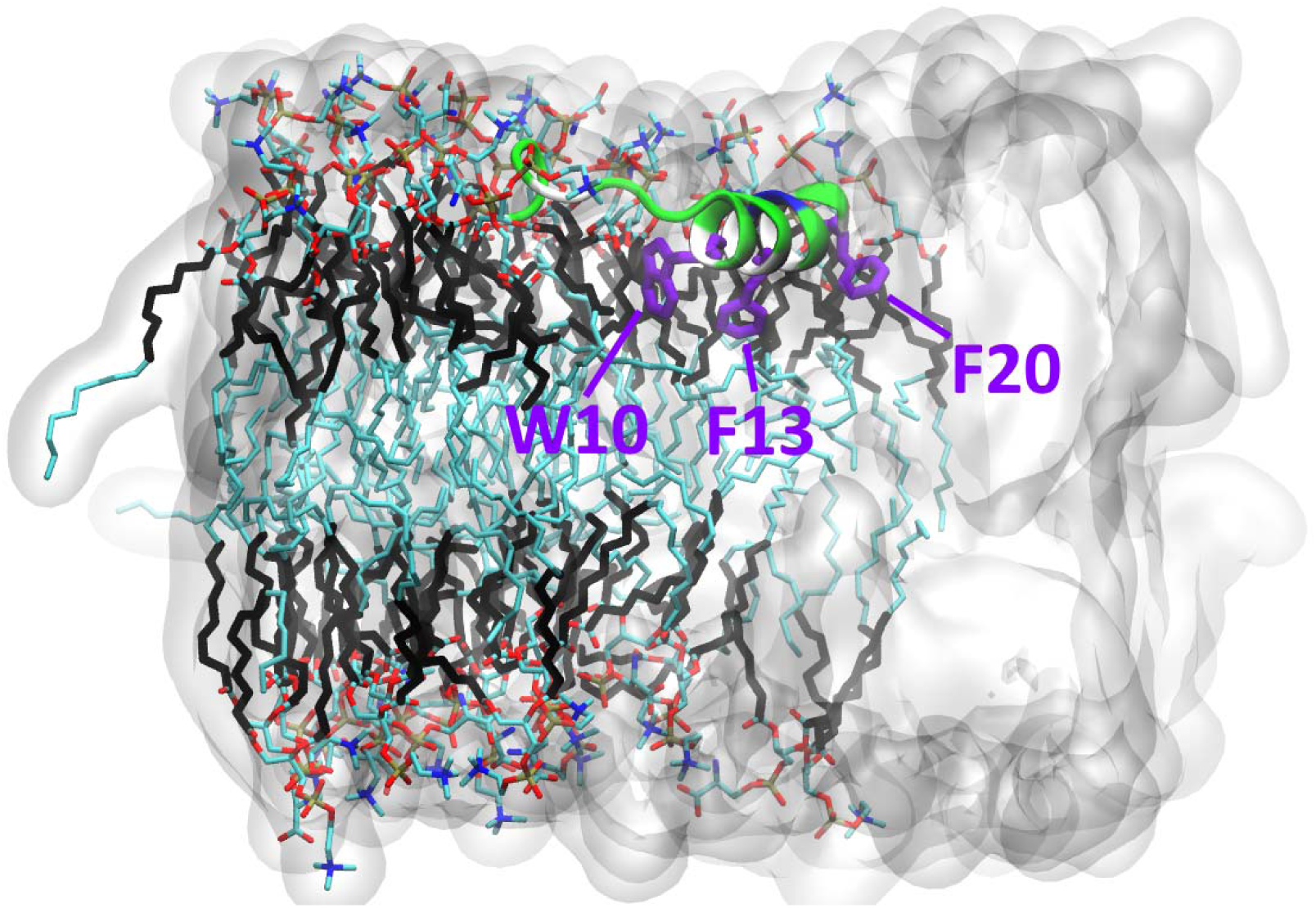
Final snapshot of the position and orientation of the tryptophan and the phenylalanine amino acids (highlighted with purple licorice representation) in the POPC-POPS (4:1) membrane bilayer. This snapshot is from the simulation represented in panel B and D of Figure 7 and shows the bilayer from a different angle than shown in Figure 7. The lipid is color-coded so that CH_2_ above the double bond is black, CH_x_ below the double bond is cyan, oxygen atoms are red, and phosphate groups are tan.

### 3.3. Binding Affinity for ALPS Peptide Binding to Membranes

We next evaluated the strength of the thermodynamic driving force for binding of the Osh4 ALPS peptide to model membranes. As noted above, tryptophan fluorescence is highly effective for studying how peptides interact with model membranes, so we also utilized tryptophan fluorescence data to estimate the binding constant for the ALPS-membrane interaction. To do this, we performed titration measurements of the ALPS peptide with 4:1 POPC-POPS vesicles (**Figure 9**). The fluorescence data were corrected for scattering artifacts using Ladokhin et al.’s correction method [61] that requires fluorescence measurements of pure tryptophan interacting with the identical model membrane under identical conditions. Using the corrected titration data and the equations described by Tatulian [62–64], we determined a dissociation constant (*K_d_*) of 1.88 µmol/L ± 0.47 µmol/L, which corresponds to a Δ*G_Fluoro_* of −32.76 kJ/mol ± 0.63 kJ/mol (−7.82 kcal/mol ± 0.15 kcal/mol) and to 7.34 ± 0.08 lipids bound per peptide molecule (*N*). The equations used by Tatulian translate raw fluorescence titration data into binding parameters by treating the lipid bilayer as a realistic, saturable surface. Specifically, these equations describe a fundamental reaction (**Equation 3**) after accounting for lipid and peptide concentrations, lipid accessibility, lipids bound per peptide, relative fluorescence intensity, and fractional saturation.

**Figure 9.**
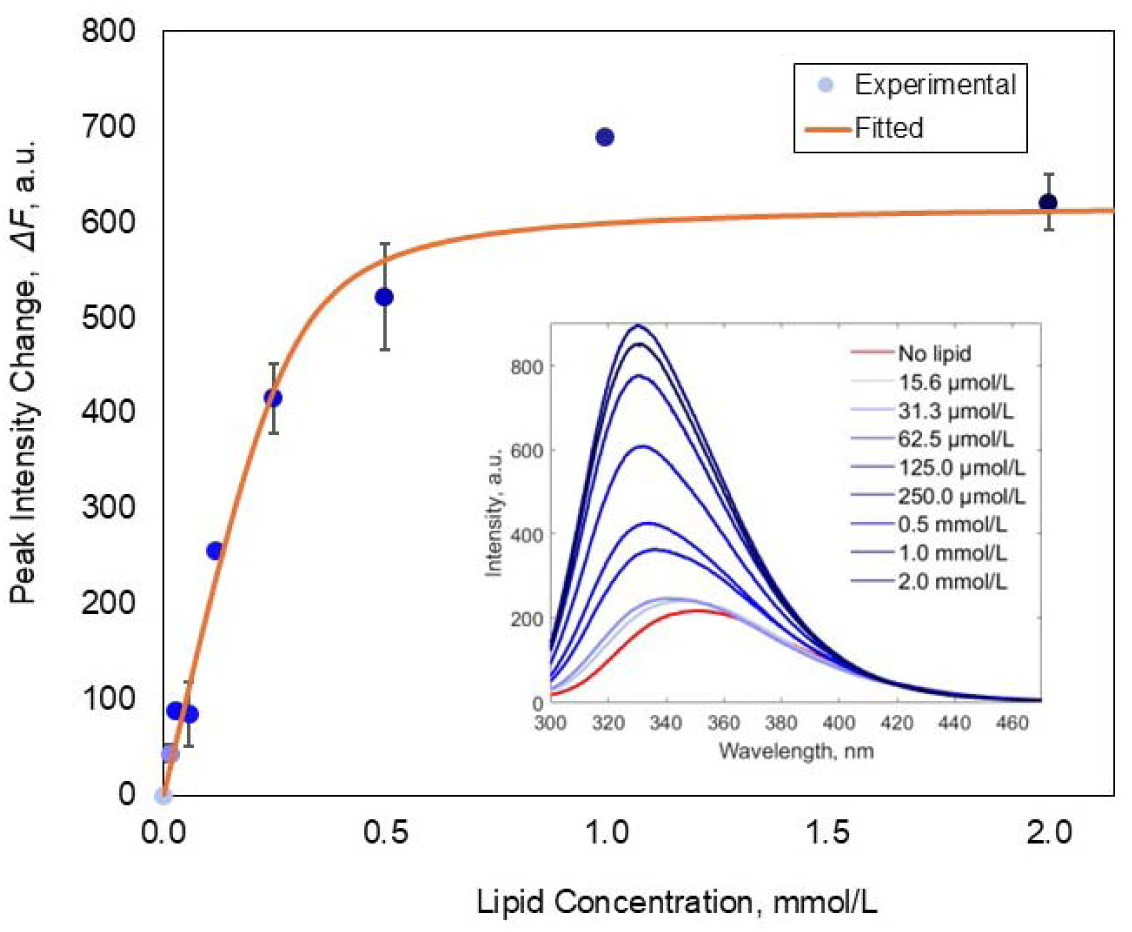
Change in fluorescence intensity resulting from the titration of 20 µmol/L ALPS peptide with 4:1 POPC-POPS LUVs at 298 K. The change at each lipid concentration is relative to the intensity of the peptide alone. The blue dots represent the corrected peak intensity changes using tryptophan zwitterion fluorescence as a reference, and the red line represents the model fit to the corrected intensities. Error bars represent the standard error of the mean (*n* = 2). The inset shows the data used to identify the peak intensity at each lipid concentration.

We also evaluated the binding of the ALPS peptide to LUVs using isothermal titration calorimetry (ITC). ITC measures binding free energy from a binding isotherm and is a more direct measurement of binding than tryptophan fluorescence, which uses data based on a single residue. The binding isotherms (**Figure 10**) were determined by injecting either POPC or 4:1 POPC-POPS lipid vesicles into an ALPS peptide solution, i.e., by gradually reducing the free peptide concentration in the reaction cell. The enthalpy of binding of the ALPS peptide to the LUV (Δ*H_ITC_*) was determined, as well as dissociation constant (*K_d_)* and apparent stoichiometry (*N)* (**Table 2**). Because lipid was the titrant and the total lipid concentration was used for all lipid-to-peptide ratios, *N* is the apparent number of lipid molecules per bound peptide, and *K_d_* is expressed per mole of total lipid. The ALPS peptide inserts into the membrane interface, and many peptides can bind a single vesicle. We therefore used the one-site model only as an empirical description of peptide partitioning into the membrane and to extract apparent affinities and enthalpies. Binding to both membranes was favorable and predominantly entropy-driven: the enthalpy change was small and negative (Δ*H*=-1.32 for 4:1 POPC-POPS and Δ*H*=-0.84 kcal/mol for POPC), while the entropic contribution was larger (*-T*Δ*S*=-5.9 kcal/mol for 4:1 POPC-POPS and *-T*Δ*S*=-6.0 kcal/mol for POPC), consistent with hydrophobic insertion of the peptide. The favorable binding for the ALPS peptide comes from a combination of the negative binding enthalpy and the positive entropy change. Fitting a single-site binding model to the ITC data shows that the ALPS peptide’s binding affinity for 4:1 POPC-POPS vesicles (*K_d_*=7.2 µmol/L) is slightly stronger than the affinity for pure POPC (*K_d_*=11.6 µmol/L) as shown in **Table 2**. For the 4:1 POPC-POPS vesicles, we determined a Δ*G_ITC_* value of −7.23 kcal/mol, which is similar to the Gibbs free energy derived from fluorescence spectroscopy titration (Δ*G_Fluoro_*), calculated to be −7.82 kcal/mol. shows the binding isotherm resulting from the integration of each injection peak. The experiments were repeated at least two times, and representative results are shown.

**Figure 10.**
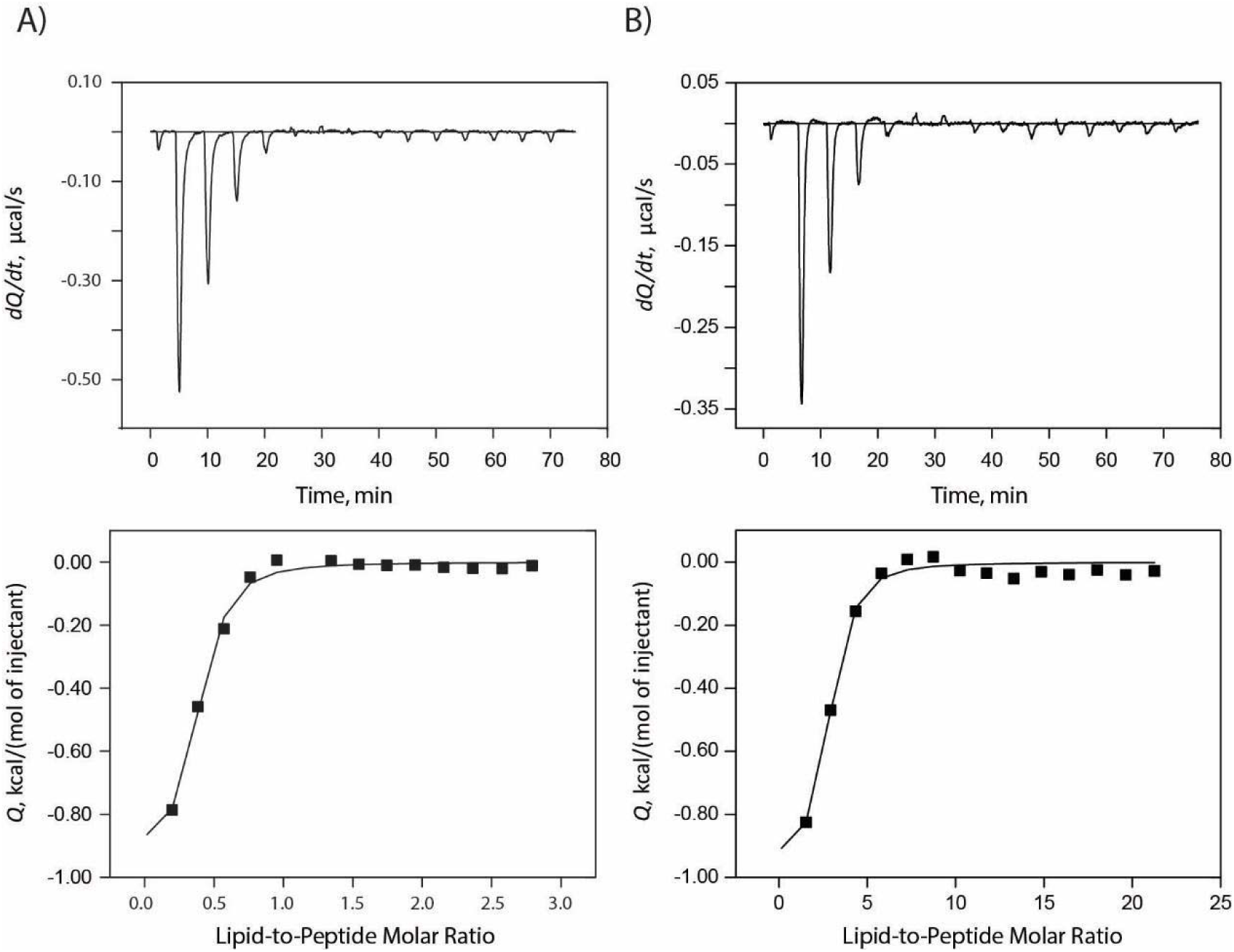
Binding of the ALPS peptide to (A) POPC vesicles and (B) 4:1 POPC-POPS vesicles. POPC LUVs (1 mmol/L) and 4:1 POPC-POPS LUVs (2 mml/L) were separately titrated into a cell containing 100 µmol/L to 150 µmol/L ALPS peptide solution. The heat (*Q*) released as a result of the interaction was measured at 298 K. The upper plot in each panel shows the thermograms generated by the series of LUV injections, while the lower plot

**Table 2.** Thermodynamic parameters^a^ obtained from ITC for the ALPS peptide binding to vesicles.

| **Membrane** | ***N*** | ***K<sub>a</sub>,***<br>**10<sup>-2</sup> L/µmol** | ***ΔG<sub>ITC</sub>,***<br>**kcal/mol** | ***K<sub>d</sub>,***<br>**µmol/L** | ***ΔH,***<br>**kcal/mol** | ***ΔS,***<br>**cal/mol/K** |
|---|---|---|---|---|---|---|
| **4:1 POPC-POPS + ALPS** | 1.62 ± 0.38 | 2.71 ± 1.25 | -7.23 ± 0.36 | 7.21 ± 4.26 | -1.32 ± 0.23 | 19.83 ± 1.95 |
| **POPC + ALPS** | 0.70 ± 0.26 | 1.32 ± 0.46 | -6.88 ± 0.28 | 11.61 ± 5.94 | -0.84 ± 0.07 | 20.23 ± 0.72 |

<sup>a</sup> Errors represent the standard error of the mean (*n* = 2).

The experimental data for the binding of the ALPS peptide to membranes allowed us to validate the accuracy of our simulated peptide-membrane interactions described by their associated force fields. To do this, we performed PMF simulations starting with the ALPS peptide bound to the full-length lipid membrane. PMF calculations can determine the energy required to pull the peptide along a realistic dissociation pathway. The starting point for each PMF simulation was the equilibrated full-length membrane simulation following conversion from the HMMM simulation. Determining the energy during the unbinding reaction requires sampling the energetic effects as the membrane and water rearrange while the peptide is slowly pulled off the membrane. By adding an external force into the simulation that pulls the peptide off the membrane, we could record the amount of force used and determine the energy profile of the unbinding reaction. The Boltzmann average of three PMF trials for Δ*G* was −38.5 kcal/mol (**Table S1**), which is significantly lower than the Δ*G* values obtained from ITC and tryptophan fluorescence data. Based on our experimental and simulation data, the protein/lipid force field (CHARMM36m/CHARMM36) overestimates the binding energy of ALPS. The force field has been shown to overestimate binding energies in the past [65–67], and a CUFIX force field [68], which was developed to address the overestimation for cationic binding, may improve the estimations. The Osh4 ALPS peptide is only mildly cationic with a single lysine. While cationic interactions are important, this peptide takes a balanced approach with multiple hydrogen bonds formed between serine and threonine amino acids and polar head groups, as well as several strong hydrophobic interactions, during the binding process. The ΔΔ*G* from ITC between ALPS with POPC and ALPS with POPC-POPS is −0.35 kcal/mol, which is low. Clearly, the simulations should indicate some energetic enhancements from electrostatic interactions with POPS, but we would need to improve the force field to better match the ITC and tryptophan fluorescence data to reproduce these findings.

We next estimated the enthalpic contribution of each amino acid to membrane binding with CHARMM interaction energy calculations. Like the calculated Δ*G*, these interaction energies are error-prone, but they enable us to assign an energy of interaction (enthalpic) to each amino acid to compare their contributions (**Figure 11**). One interesting observation is that the unfolded section in the first 10 amino acids, which is rich in serine, has stronger interaction energy on average compared to the more helical section in the later 10 amino acids (**Figure 11**). The two main outliers are the K15 and F20 residues. The salt bridge formed with K15 and the membrane is likely too strong in the CHARMM36m/CHARMM36 force field, and adjustments like CUFIX might be needed to reduce the attraction of the ALPS peptide to the model membranes. Strong contributions to the binding from both the polar amino acids (serine and threonine) as well as the bulky hydrophobic amino acids including W10 and F20 are apparent. However, F13 did not play a major role in membrane interaction but, as noted previously, could be important for π−π stacking to maintain helical stability.

**Figure 11.**
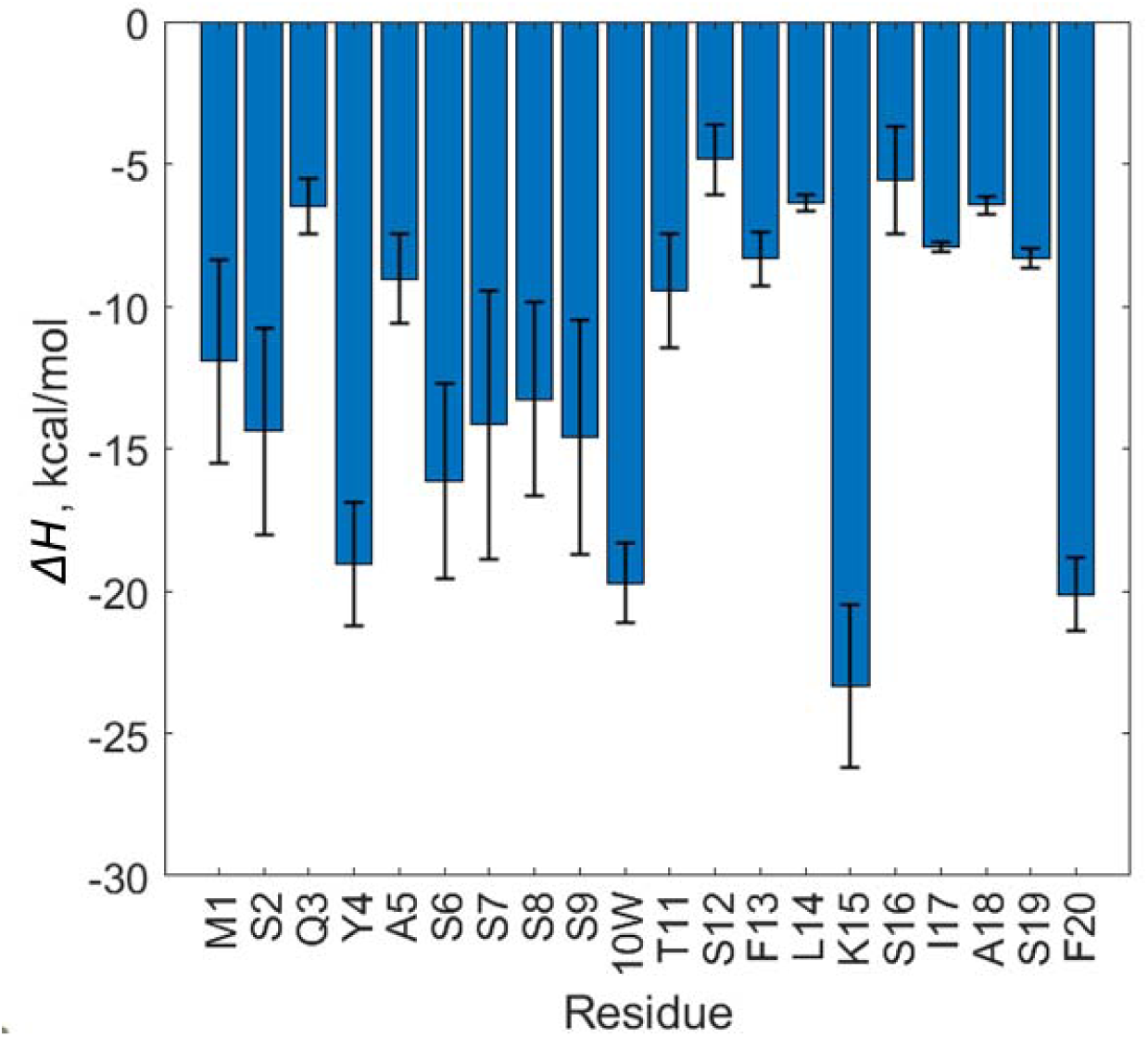
Interaction enthalpies (Δ*H*) between each of the amino acids in the ALPS peptide and the 4:1 POPC-POPS membrane from CHARMM interaction energy calculations. The standard errors are shown for the average of three simulation trials. More detailed data with individual energies per simulation and per amino acid with binned standard error are provided in **Figure S6**.

The serine and tyrosine amino acids are perhaps the most interesting amino acids in the Osh4 ALPS peptide, not only due to their known role in characterizing curvature-sensing but also because they have interesting entropic properties which might even be related to curvature-sensing properties [18, 21, 69]. Some mutational studies have confirmed these residues are important, such as showing that these amino acids increase selectivity for a smaller liposome radius or higher curvature [18]. However, no studies have proposed a physically based mechanism explaining the role of serine and tyrosine amino acids. The bilayer morphology and the shape of the peptide combine to impact the placement of serine amino acids in the ALPS peptide, and the serine amino acids can simultaneously interact with multiple chemically distinct lipid atoms (**Figure S7** and **S8**). Figure S8 shows the frequency of contacts of different amino acids with different chemical moieties of the bilayer to get an idea of the varied chemical interactions. The polar amino acids S2, Y4, and S7 interact frequently in all layers of the membrane **(Figure S8)**. We found that these amino acids are thermodynamically stabilized at the interface of the bilayer because of their entropy due to desolvation (**Figure S7** and **S8)**. Over time, the peptide diffuses over varied membrane configurations where the head group is closer or further away from the serine, and the serine can flip up or down to burrow into the hydrophobic core or reach up into the wetted head group. The main entropic contribution of buried serine and threonine is due to the liberated water from the OH solvation shell. However, the simulations indicate that, by having contacts with both the hydrated headgroups and the de-wetted hydrophobic core, these amino acids contribute to disorder, while the lysine predictably binds to the charged lipid head group and behaves more like an ordering, rigid structure. The serine, threonine, and glutamine amino acids are stabilized by diverse chemical interactions in the bilayer interface, where the combination of diffusion along the membrane surface and the entropic reconfigurations enable balanced interactions in both the hydrophobic and hydrophilic environments [69].

We calculated the fraction occupancy of each amino acid to show the fraction of time an amino acid had contact with either the choline, phosphate, glycerol, C=O, or tail Cxx, which is defined as the contacts selected by the first 8 carbons below the unsaturation on the Sn1 and Sn2 chains (**Figure S8**). The interactions of tryptophan and phenylalanine with the membrane are relatively predictable. Based on our data, they prefer to interact in the hydrophobic core of the membrane as generally expected. The lysine amino acid exhibits unexpected behavior due to its lack of a dominant fraction occupancy in any level of the peripheral membrane. One would typically expect this amino acid to interact with the charged head group, but instead it interacts at all levels of the membrane. This behavior might arise from the fact that when embedded in the peripheral membrane, the linear aliphatic chain must interact with the hydrophobic core, while the ammonium group should simultaneously interact with the head groups.

### 3.4. ALPS Peptide Binding Influences Membrane Structure

The results described so far provide insight into the structure of the ALPS peptide during interactions with the membrane. To learn more about the effect of the peptide on membrane structure, we employed X-ray diffraction to investigate the effects of incorporating the ALPS peptide into lipid multilayers. X-ray diffraction provides the distribution of electron density across the membrane thickness to reveal the positions of the headgroups and hydrocarbon core. We formed lipid multilayers from POPC and binary mixtures of POPC and POPS at a peptide-to-lipid (P/L) molar ratio of 1:25. At this P/L, the concentration of ALPS at the bilayer surface is fairly high but less than what would be needed for complete surface coverage, which would require a bulk P/L corresponding to the value of 1/*N* where *N* is the number of lipids per bound peptide molecule. The simulation suggests average and standard error values of 8.46 ± 0.02 lipids per peptide for our three simulations (**Figure S9**), which agree reasonably well with the value of *N*=7.3 in the tryptophan fluorescence binding isotherm. Despite the high P/L used in the experiments, we determined that the repeat spacing (bilayer thickness + water layer thickness, *d*) of the multilayer system shows only a modest decrease in the presence of ALPS, compared to peptide-free POPC or POPC-POPS bilayers (**Figure 12A**, **Table 3**). The most pronounced thinning (2.17 Å) was observed in 1:1 POPC-POPS multilayers, consistent with the observation that the peptide partitions more strongly in bilayers containing POPS. The significant increase in electron density in the headgroup regions of bilayers with ALPS, compared to neat bilayers, indicates that the peptide adopts an interfacial location at the bilayer-water interfaces. The intercalation of the ALPS peptide between the lipid headgroups causes a bilayer area expansion at constant hydrocarbon density, which is commonly observed for peptides known to interact with membranes [70–72]. Similar to the results for *d*, we also observed a thinning trend for the headgroup-to-headgroup distance (*d_HH_*), which shows a maximal decrease with ALPS in 1:1 POPC-POPS multilayers (**Table 3**). The thinning effects were accompanied by broadening (loss of structural features) in the hydrocarbon region of the bilayer. This can be explained by an increased disorder of the acyl chains, which is normally associated with bilayer area expansion. Overall, the perturbations brought by ALPS to the bilayer structure (Figure 8) indicate a substantial peripheral binding of the peptide and insertion into the hydrophobic core, though the X-ray scattering profile is minimally perturbed. Binding is more pronounced when an anionic lipid such as POPS is present.

**Figure 12.**
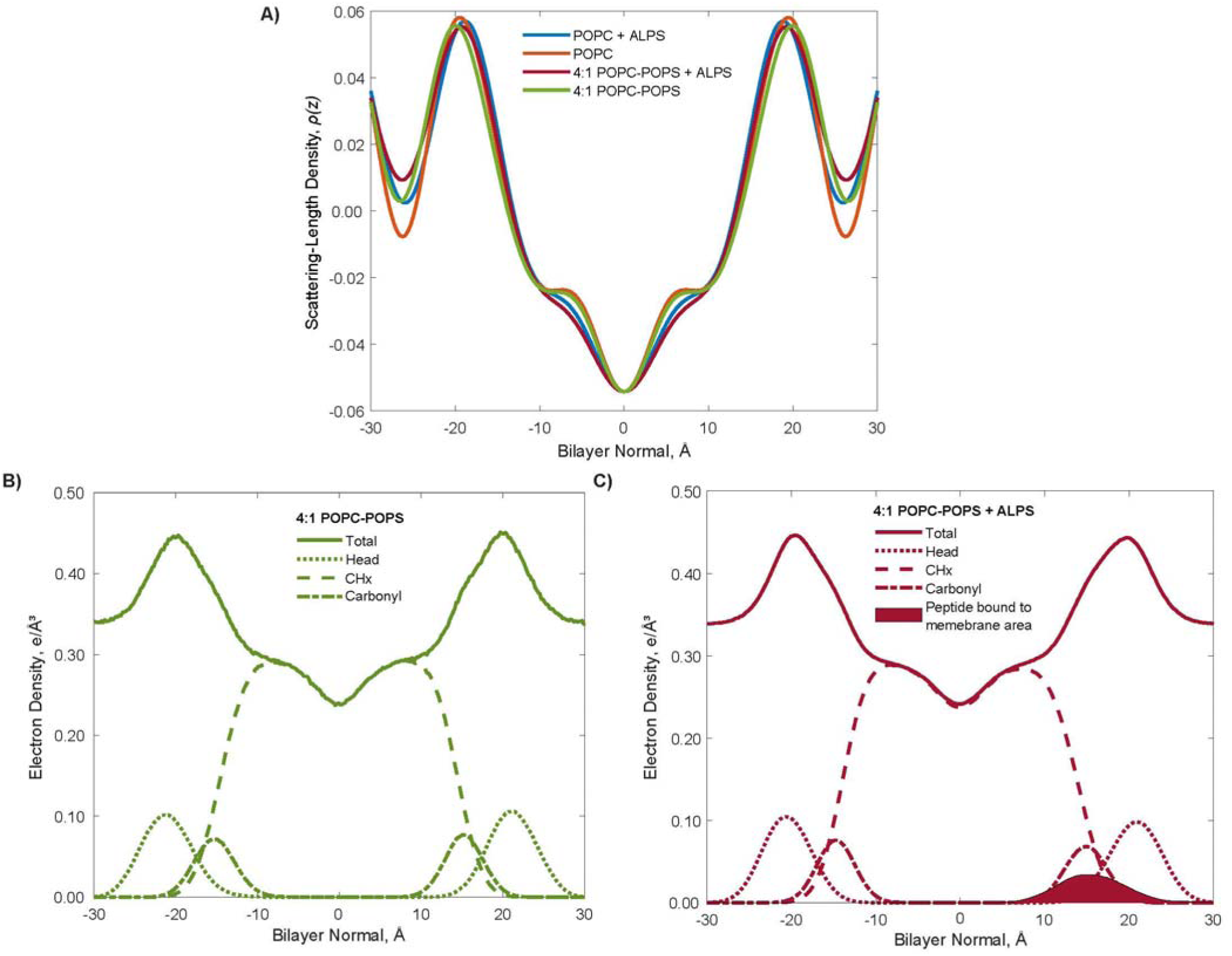
Electron density profiles for experimental and simulated systems of membranes and ALPS peptide. **(A)** X-ray diffraction scattering-length density profile of pure POPC and 4:1 POPC-POPS bilayers alone and in the presence of the ALPS peptide at a peptide-to-lipid molar ratio of 1:25 and at 97 % relative humidity measured at 298 K. The bilayer normal distance represents the distance from the center of the hydrophobic core of the lipid bilayer. (B) MD-derived electron density profiles for the ALPS peptide binding to a 4:1 POPC-POPS bilayer with no peptide. **(C)** MD-derived electron density profiles for the ALPS peptide binding to a 4:1 POPC-POPS bilayer with peptide. For the MD-derived profiles, the overall profile, along with the contributions from the lipid head group, CHx groups, and carbonyl groups are shown.

**Table 3.** The membrane parameters^a^ obtained from X-ray diffraction measured in the presence and absence of ALPS peptide at a peptide-to-lipid molar ratio of 1:25 measured at 298 K.

| Membrane | ***d*, Å** <sup>b</sup> | ***Δd*, Å** | ***d<sub>HH</sub>*, Å** <sup>b</sup> | ***Δd<sub>HH</sub>*, Å** |
|---|---|---|---|---|
| **POPC** | 52.45 ± 0.13 | 0.61 | 38.92 ± 0.02 | 1.24 |
| **POPC + ALPS** | 51.84 ± 0.17 |  | 37.68 ± 0.10 |  |
| **9:1 POPC-POPS** | 52.98 ± 0.13 | 0.45 | 39.16 ± 0.03 | 0.64 |
| **9:1 POPC-POPS + ALPS** | 52.53 ± 0.28 |  | 38.52 ± 0.08 |  |
| **4:1 POPC-POPS** | 53.02 ± 0.08 | 0.46 | 39.88 ± 0.04 | 1.28 |
| **4:1 POPC-POPS + ALPS** | 52.56 ± 0.21 |  | 38.60 ± 0.12 |  |
| **1:1 POPC-POPS** | 54.89 ± 0.15 | 2.17 | 41.05 ± 0.05 | 1.65 |
| **1:1 POPC-POPS + ALPS** | 52.72 ± 0.26 |  | 39.40 ± 0.05 |  |

**<sup>a</sup>** *d* represents the repeat spacing (bilayer thickness plus water layer thickness), *Δd* is the change in repeat spacing, *d<sub>HH</sub>* is the headgroup-to-headgroup distance, and *Δd<sub>HH</sub>* is the change in headgroup-to-headgroup distance.
**<sup>b</sup>** Error represents one standard deviation.

We can compare the membrane density obtained from simulations to the experimental results. Even though the peptide is more diluted in the simulation than in the experiment, the simulations provide an atom-level analysis of the interacting peptide-bilayer system that can aid in interpreting the experimental data. The MD-derived electron density profiles of the four simulated systems show in detail where the peptide is localized with respect to the other membrane components (**Figure 12B** and **12C**). From our simulations of the electron density profile, the peptide widened and lowered the head group peak, yielding a more diffuse head group-water interface, but the most prominent membrane change occurs in the distance between the half maximum of the CH (hydrocarbon) profile, 2*D_c_*, with a 1.1 Å decrease upon the addition of the peptide (**Table 4**). Twice the distance from the center to where the water density reaches its half-maximum, *D_B_*, also decreased with the introduction of the peptide, but to a smaller extent (0.5 Å). The head-to-head distance, *D_H_*, is the distance between the overall peak intensity, usually the distance in the electron density profile between the phosphates from the two leaflets. While the values of 2*D_c_* from the simulations and *d_HH_* from the X-ray diffraction data differ, the simulation Δ(2*D_C_*) matches the qualitative trend in the experimental Δ*d_HH_*, with both indicating a thinning of the membrane in the presence of the peptide. *D_H_* is most closely related to the *d_HH_* in the measured data, but the simulation shows little peptide binding effect for *D_H_*. These differences could be caused in part by the low hydration of the experiment and different P/L ratios used in the measured versus simulated experiments. In any case the thinning caused by the ALPS peptide is minor compared to that caused by antimicrobial peptides at similar P/L ratios, indicating that the ALPS peptide does not perturb the bilayers significantly.

**Table 4.** Changes in the distance of the overall peak intensities, CH hydrocarbon, and water based on the electron density profiles for MD simulation of ALPS binding to a 4:1 POPC-POPS bilayer.

| **Membrane** | **2*D<sub>C</sub>*, Å<sup>a</sup>** | ***Δ*(2*D<sub>C</sub>*), Å** | ***D<sub>B</sub>*, Å** <sup>c</sup> | ***ΔD<sub>B</sub>*, Å** | ***D<sub>H</sub>*, Å<sup>b</sup>** | ***ΔD<sub>H</sub>*, Å** |
|---|---|---|---|---|---|---|
| **4:1 POPC-POPS** | 29.0 | 1.1 | 39 | 0.5 | 40 | 0.2 |
| **4:1 POPC-POPS + ALPS** | 27.9 ± 0.1 |  | 38.5 ± 0.1 |  | 39.8 |  |

<sup>a</sup> Distance between the half maximum of the CH (hydrocarbon) profile. Error represents the standard error of the mean (*n* = 3 for the simulation with peptide and *n*=1 for the simulation without peptide).
<sup>b</sup> Distance between the overall peak intensity of the electron density profile from the two leaflets. Error represents the standard error of the mean (*n* = 3 for the simulation with peptide and *n*=1 for the simulation without peptide).
<sup>c</sup> Position of the half maximum of the water intensity. Error represents the standard error of the mean (*n* = 3 for the simulation with peptide and *n*=1 for the simulation without peptide).

## 4. Conclusions

We combined experimental data obtained using circular dichroism, fluorescence, ITC, and X-ray scattering with MD simulations to gain insight into the binding of the Osh4 ALPS peptide to membranes. The Osh4 protein fits into a broader context of lipid transport occurring at the TGN-ER interface and is one of many proteins responsible for maintaining the concentrations of lipids like PI(4)P [11, 12, 31, 35]. The ALPS-like motif of Osh4 is important in the lipid transport function of this protein but the molecular-level details of its structure and binding characteristics to membranes are less known. The peptide structure changes from a mixture of unfolded and partially α-helical peptide in the aqueous phase to a structure dominated by α-helical content in the membrane, as observed from CD measurements. The helical fraction was shown to be directly correlated with the amount of anionic lipids in the membrane. We found that the aqueous phase T-REMD simulations slightly over-stabilized the α-helical structure in water in comparison to the CD results, which might be caused by stronger than expected π−π stacking in the CHARMM force field.

Measuring the free energies of peptide binding to determine enthalpic and entropic contributions to binding is a highlight of this work. ITC measures the total thermal reaction, including heat from binding and folding. We obtained similar binding constants from ITC and fluorescence spectroscopy for binding of the ALPS peptide to 4:1 POPC-POPS. Our ITC data suggest the enthalpy change was favorable, and ALPS binding is overall an entropy-driven process with nearly six-fold more energy coming from entropy than from enthalpy. The major entropic part of the binding could come from desolvation of the membrane and peptide, and investigating the serine, threonine, and glycine residues and their entropic contributions to the membrane could yield additional insight.

Comparison of the experimental results with the simulation results indicates significant room for improvement in the force field interactions between membranes and peptides like the ALPS peptide. Specifically, simulations suggest a five-fold tighter binding free energy than those measured from ITC and tryptophan fluorescence. CUFIX would improve electrostatic inaccuracies involving the lysine residue, but other challenges include accurate π-π stacking contributions and the importance of secondary structure formation. Small-molecule ITC could be used to model the interaction of lipids and amino acids to refine the force field in the future and allow simulations to provide a more quantitatively accurate description of peptide-lipid interactions.

We used X-ray diffraction studies to measure changes in the membrane structure due to peptide binding, and our results indicate a greater thinning of the phospholipid bilayer with the presence of POPS lipids. The experimental results were supported by simulations; both sets of results indicate only a minor perturbation of the membrane structure due to the binding of the ALPS peptide and no major reorganization.

Our results emphasize the selectivity of the Osh4 ALPS motif for specific lipid environments. This selectivity could contribute to Osh4’s role in moving lipids between membranes. Since the TGN is known to have more anionic lipids compared to the nearly negligible amount in the ER, it is likely that the ALPS-motif would preferentially bind to the TGN compared to ER. Our ITC measurements demonstrate a higher affinity to POPC:POPS membranes albeit by only a 0.4 kcal/mol difference. However, it is expected K15-lipid interactions would promote faster kinetics in binding to TGN compared to ER without anionic lipids. Our work also demonstrates that differential binding to lipids affects the motif’s ability to induce conformational changes in the membrane, which plays a critical role in intracellular lipid trafficking, membrane organization, and overall targeting in the development of Osh4’s ability to form MCSs.

## 5. Acknowledgments

We thank Dr. Taylor Woehl for access to his lab’s fluorometer. We thank Sijie Hao from the University of Maryland BioWorkshop for training on circular dichroism. In part, this work was supported by the National Science Foundation (MCB 1951425), the National Institutes of Health (R35GM161807), and a University of Maryland Summer Research Fellowship.

All simulations were conducted using the Extreme Science and Engineering Discovery Environment (XSEDE) which are supported by NSF grant ACI-1548562 and the NSF Office of Advanced Cyberinfrastructure awards #2138259, #2138286, #2138307, #2137603, and #2138296 under the MCB100139 allocation. The Maryland Advanced Research Computing Center (MARCC) supercomputers including Deepthought2 and Zaratan. Data analysis was carried out on the MARCC supercomputer and the University of Maryland’s Zaratan, which is managed by the Division of Information Technology.

## Disclaimer

Certain commercial equipment, instruments, or materials are identified in this paper in order to specify the experimental procedure adequately. Such identification is not intended to imply recommendation or endorsement by the National Institute of Standards and Technology, nor is it intended to imply that the materials or equipment identified are necessarily the best available for the purpose. Official contribution of the National Institute of Standards and Technology; not subject to copyright in the United States.

## Data sharing statement

The data that support the findings of this study are available from the corresponding authors upon reasonable request.

