## Supporting Figures and Tables for "Elucidating the membrane-binding mechanism of the ALPS-like motif of the Osh4 protein"

**Table S1.** ΔG for the three replicates of the ALPS peptide with membranes composed of 80 % POPC and 20 % POPS determined in MD simulations. The Boltzmann average weights the lowest free energy more than other configurations. Other metrics include the comparison with the wet-lab value, the area/lipid, and the % helical content. The standard errors for the free energies are based on the three replicates. The standard error for the area/lipid is for the last 100 ns of the simulation.

| Membrane | Trial | ΔG, kcal/mol | Boltzmann Average, kcal/mol | Wet-Lab, (kcal/mol) | Area/Lipid | % Helical |
|---|---|---|---|---|---|---|
| | 1 | -22.8 ± 4.6 | | | 66.6 ± 0.2 | 50 |
| 4:1 POPC-POPS | 2 | -38.5 ± 4.6 | -38.5 ± 4.6 | -7.6 | 66.0 ± 0.1 | 55 |
| | 3 | -28.7 ± 4.6 | | | 66.0 ± 0.2 | 60 |

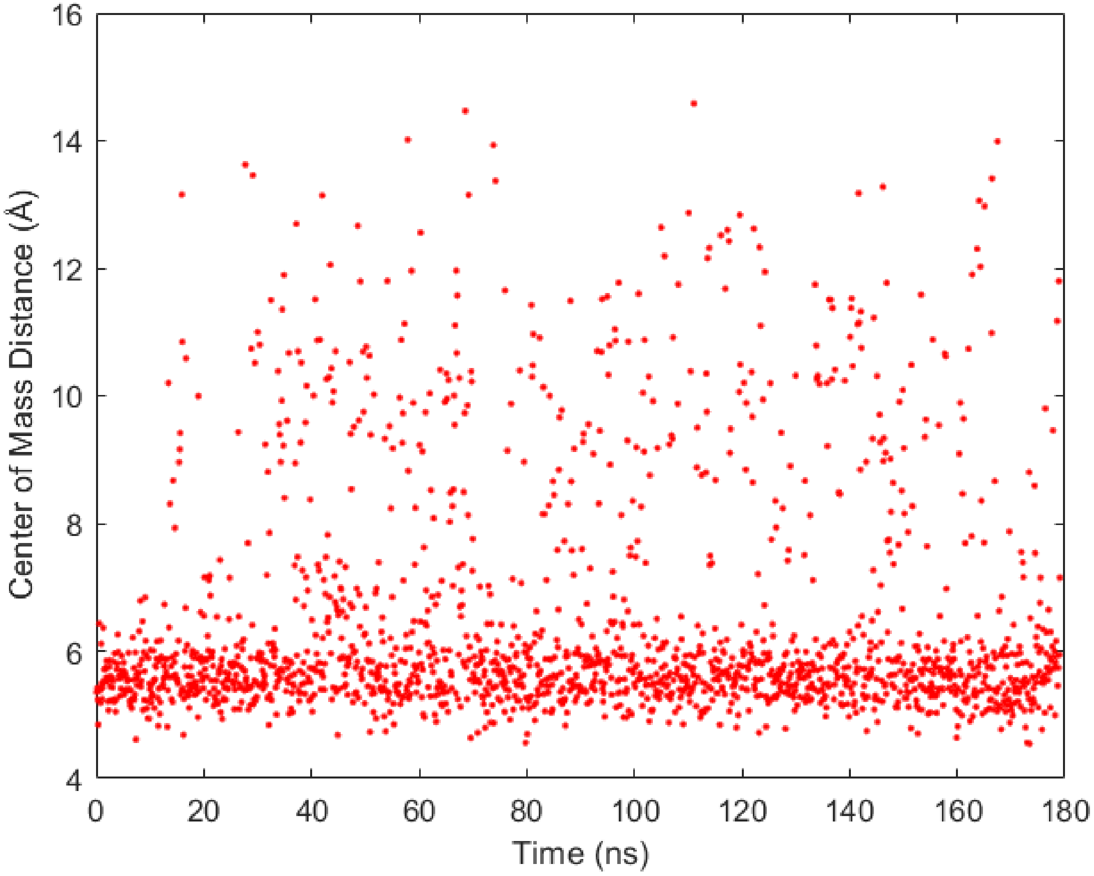


**Figure S1.** Distance between W10 and F13 of the ALPS peptide over time in MD simulations. The cutoff for $\pi$-$\pi$ is not clearly documented for the CHARMM force field, but these data suggest that the average value is between 5 Å to 6 Å.

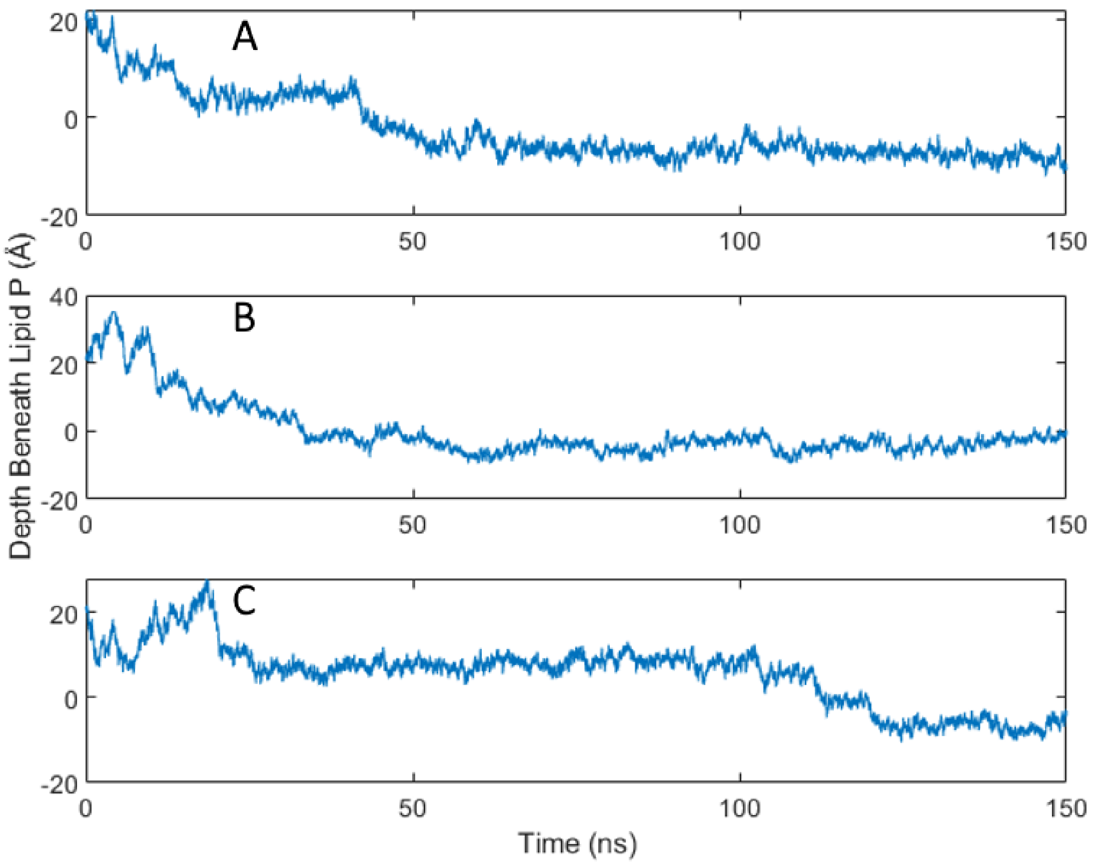


**Figure S2.** Distance between the ALPS peptide and the lipid phosphate over time. Each panel shows data for one of three replicates of the HMMM model. The distance was calculated as the distance between the average position of the alpha carbon carbons in the peptide and the position of the phosphate of the leaflet the peptide was associated with. The 0 depth represents the location center of mass of the lipid phosphate.

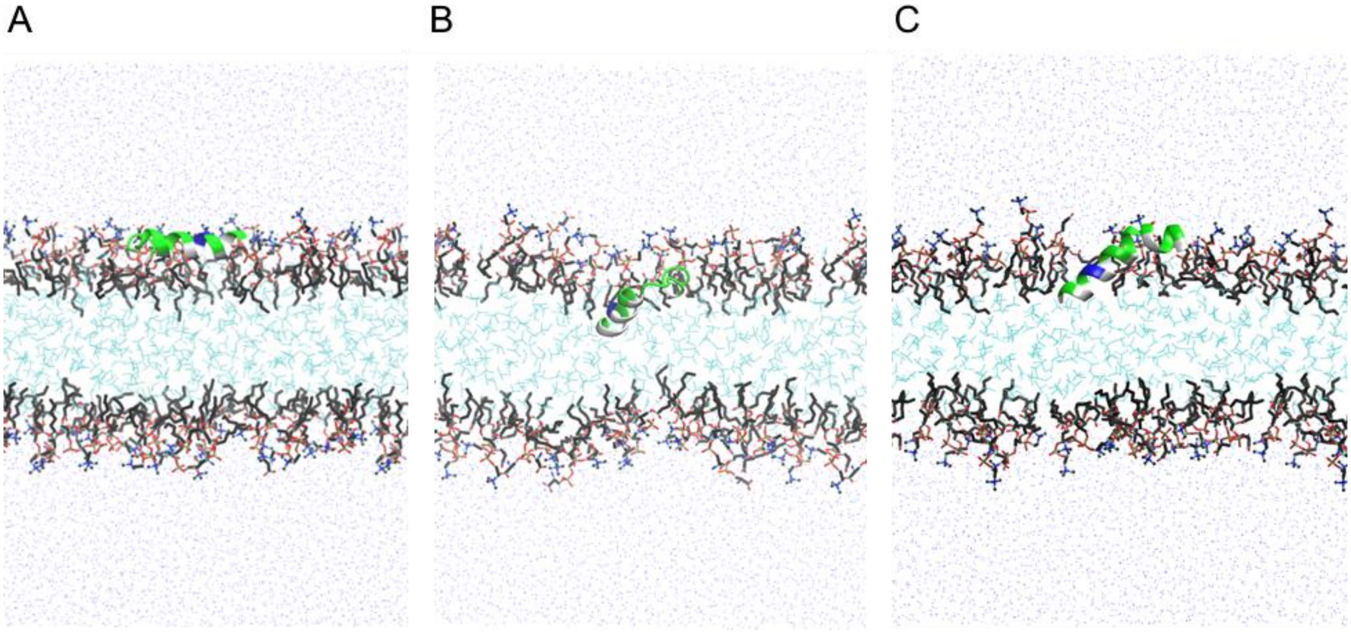


**Figure S3.** HMMM simulations of the Osh4 ALPS peptide bound to a membrane with 80 % POPC and 20 % POPS. Each panel (A-C) represents a distinct simulation. The simulations were run for 150 ns in HMMM.

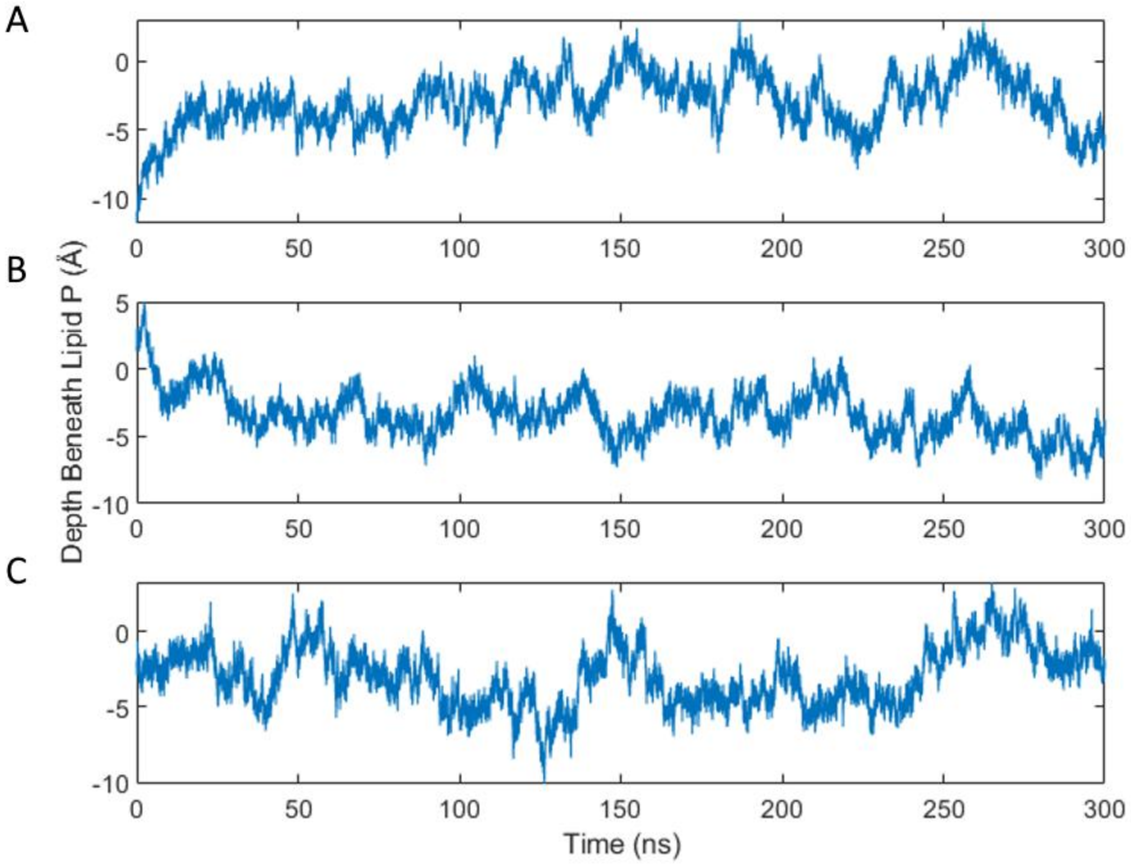


**Figure S4.** Time courses of the insertion depths of the alpha carbons of the $10^{th}$ through $20^{th}$ residues of the peptide simulated in a full-length lipid bilayer (80 mol% POPC and 20 mol% POPS). Each panel represents one replicate. The penetration distances are measured beneath the level of membrane phosphates.

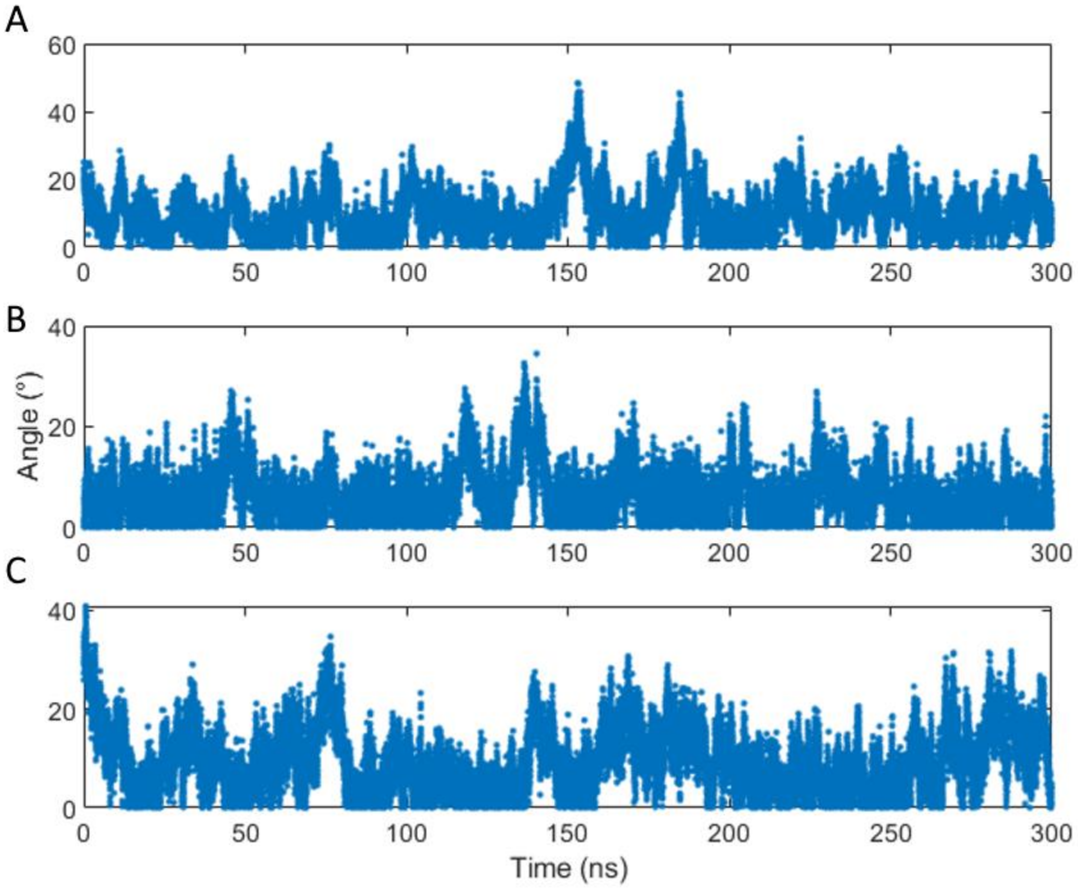


**Figure S5.** Angular position of the 10th through 20th residue segment of the ALPS peptide in POPC-POPS membrane bilayers simulated in full-length lipids. Each panel shows a separate replicate simulation. The angle is the angle between the membrane normal and the helix formed by the 10th through 20th amino acids.

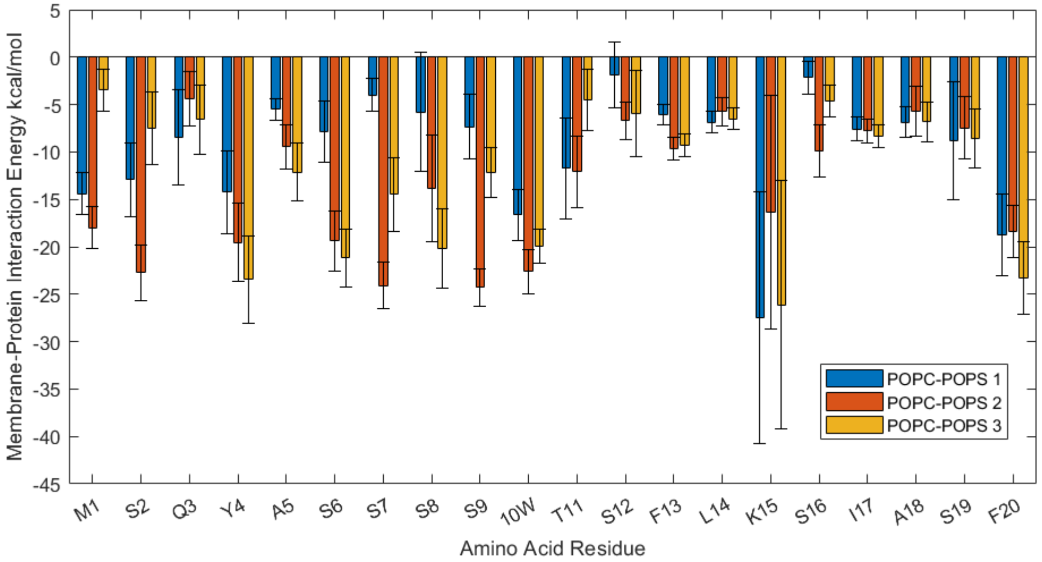


**Figure S6**. Interaction energy between the membrane and the peptide calculated from 20 binned averages over the course of 28 ns. Error bars represent standard error.

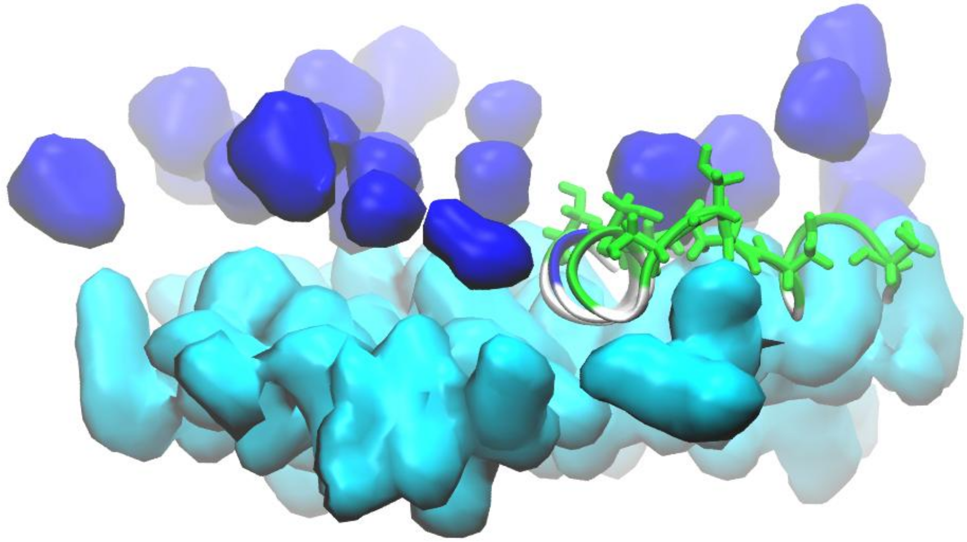

**Figure S7.** Interaction of the Osh4 ALPS peptide with a membrane (80 mol % POPC and 20 mol % POPS), illustrating the importance of the serine amino acid in the amphipathic nature of the peptide. The serine amino acids in the ALPS peptide are shown in a green licorice representation. The membrane hydrophobic core is shown in cyan, and the choline groups are shown in blue surface representations; these groups define the upper and lower limits of the peripheral membrane. The hydrophobic tryptophan and valine residues anchor the peptide in the membrane. The serine amino acids appear to have an affinity for both the polar head and parts of the hydrophobic core.

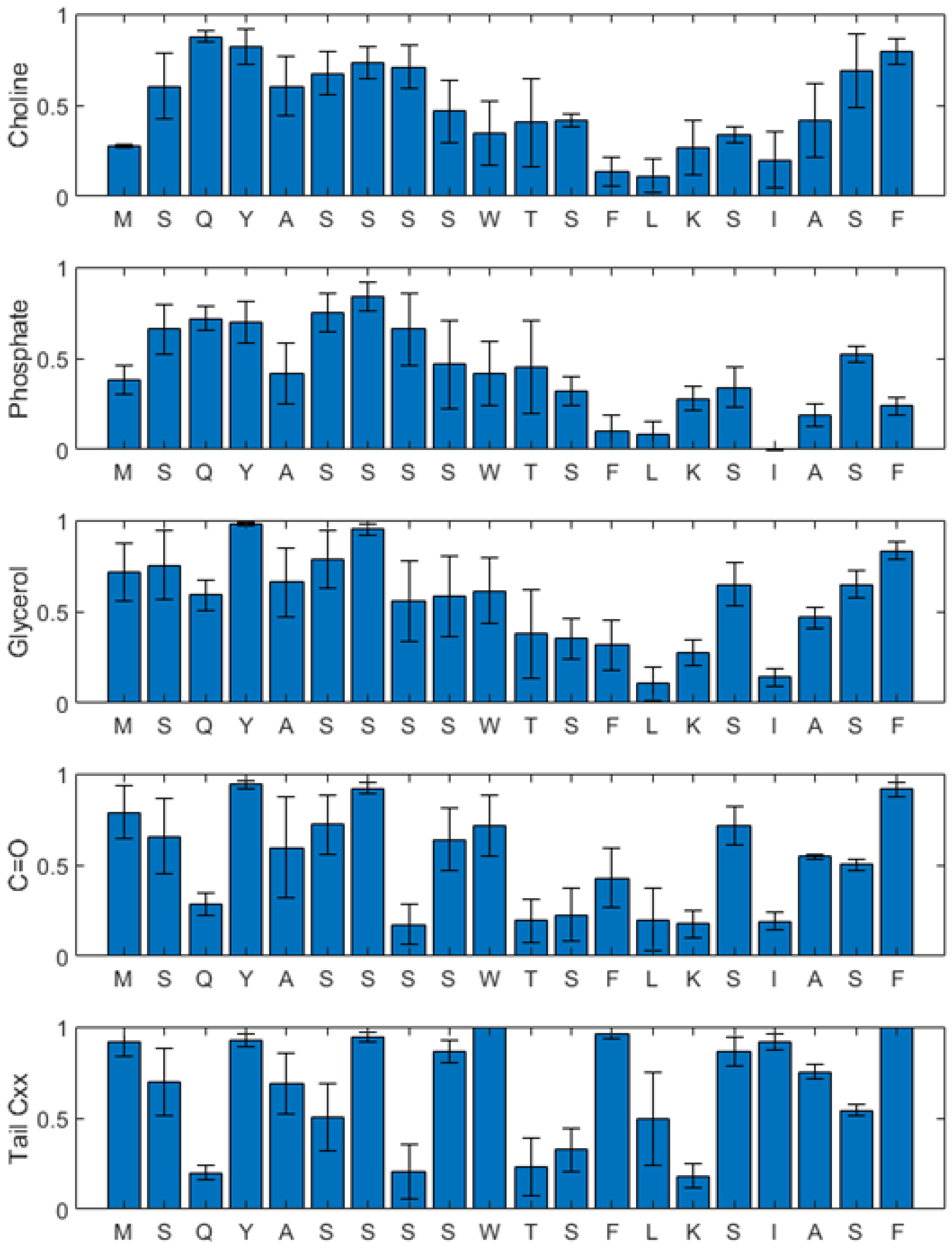


**Figure S8.** Average number of contacts between ALPS peptide amino acids and POPC lipid atoms or groups of atoms. Contacts were informed by the unbiased dynamics generated from the force field using a 4.5 Å cutoff to indicate contact. The number of contacts was calculated in every other frame in the last 126 ns of the trajectory (12600 frames). Error bars represent standard error.

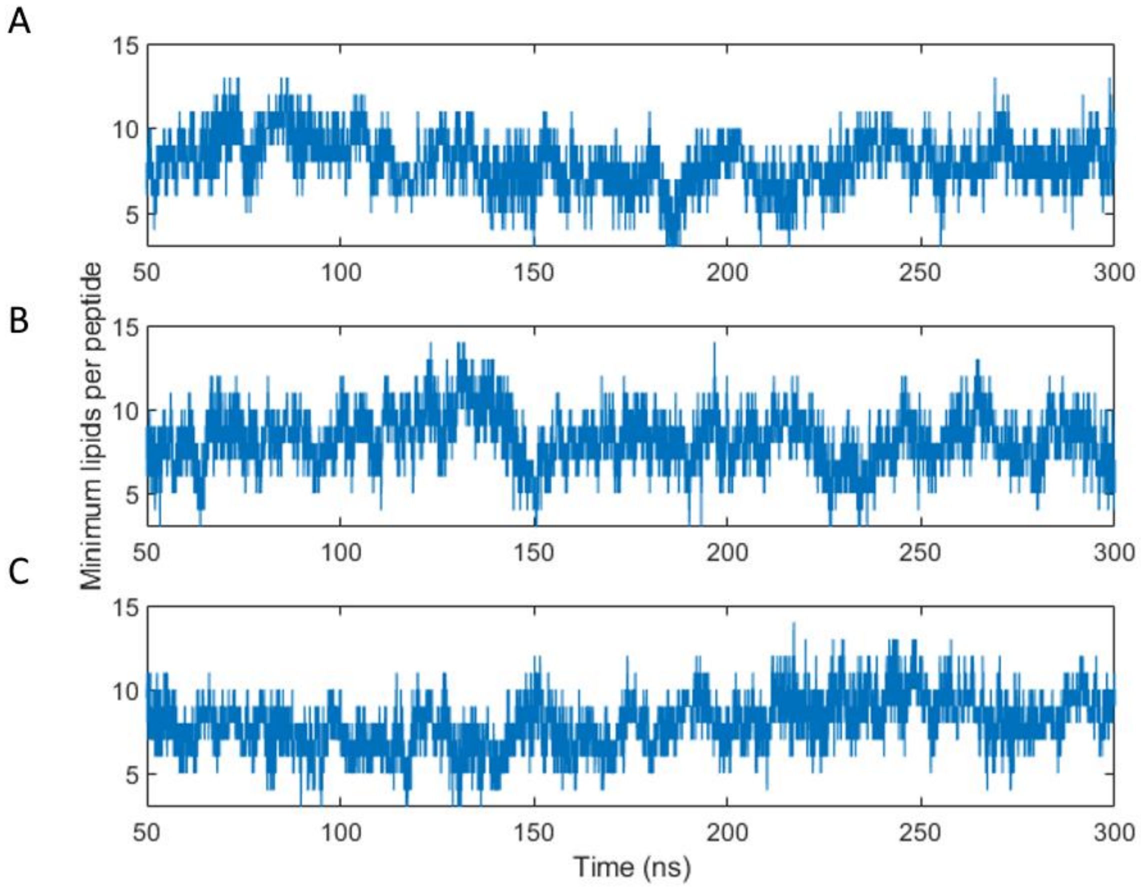


**Figure S9.** Number of lipids bound per ALPS peptide molecule for the 80 mol % POPC and 20 mol % POPS bilayers. Each panel corresponds to a separate replicate and plots the number of unique phosphorus atoms within 6 Å of the peptide over the course of a 300 ns simulation. These results show how the value of N determined by ITC and fluorescence spectroscopy is related to physical thermodynamic properties that fluctuate over time.

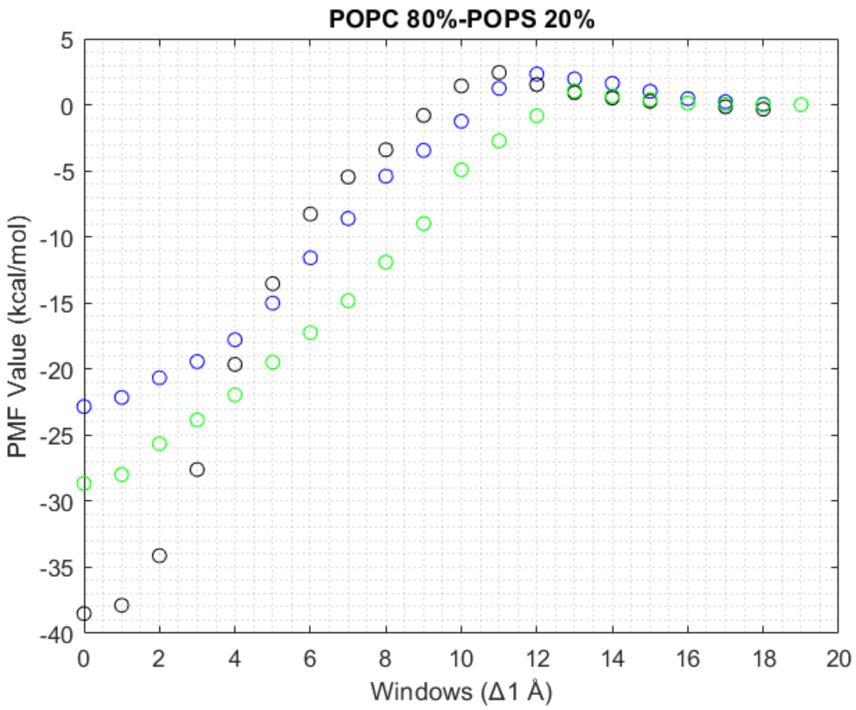


**Figure S10.** PMF profile for the ALPS peptide and a membrane with 80% POPC and 20% POPS lipids. The three sets of data represent the three replicate PMF trials, and the final free energy differences correspond to the values found in Table S1. The profile shows the energy value of each 1 Å window in the full reaction profile, where each window is a separate simulation with a different collective variable set point. The same information can be used to construct a probability distribution showing how likely the peptide could be found in different parts of the membrane. The boot strapped error is smaller than the circles used to plot the data.
